# Stability of Genomic Imprinting and X-Chromosome Inactivation in the Aging Brain

**DOI:** 10.1101/2023.09.29.560184

**Authors:** Samantha Mancino, Janith Seneviratne, Annalisa Mupo, Felix Krueger, David Oxley, Melanie A Eckersley-Maslin, Simão Teixeira da Rocha

## Abstract

Epigenetic drift is a hallmark of aging that contributes to the irreversible decline in organismal fitness ultimately leading to aging-related diseases. Epigenetic modifications regulate the cellular memory of the epigenetic processes of genomic imprinting and X-chromosome inactivation to ensure monoallelic expression of imprinted and X-linked genes. Whether epigenetic drift affects maintenance of genomic imprinting and X-chromosome inactivation has not been comprehensively studied. Here, we investigate the allele-specific transcriptional and epigenetic signatures of the aging brain, by comparing juvenile and old hybrid mice obtained from C57BL/6J (BL6) & CAST/EiJ (CAST) reciprocal crosses, with an emphasis on the hippocampus (HCP). We confirm that the aged HCP shows an increase of DNA hydroxymethylation, a sign of epigenetic drift, and a typical aging transcriptional signature. Genomic imprinting was found to be largely unaffected with stable parent-of-origin-specific DNA methylation in HCP, but also other brain regions such as the cerebellum (CB), nucleus accumbens, hypothalamus and prefrontal cortex. Consistently, transcriptomics analysis confirmed unaltered imprinting expression in the aged HCP. An exception are three novel non-coding transcripts (*B230209E15Rik*, *Ube2nl* and *A330076H08Rik*) at the Prader-Willi syndrome/Angelman syndrome (PWS/AS) imprinted locus which lose strict monoallelic expression during aging. Like imprinting, X-chromosome inactivation was remarkably stable with no signs of aging-driven skewing or relaxation of monoallelic expression of X-linked genes. Our study provides a valuable resource for evaluating monoallelic expression in the aging brain and reveals that, despite epigenetic drift during aging, genomic imprinting and X-chromosome inactivation remain predominantly stable throughout the process of physiological aging in the mouse brain.

## Introduction

Aging is defined as an irreversible loss of physiological integrity associated with functional decline of tissues and organs, progressively leading to aging-related illnesses, such as neurodegenerative diseases^1^. At the molecular and cellular levels, several hallmarks have been associated with aging, including epigenetic drift^2,3^. Amongst the aging-related epigenetic changes there is DNA methylation, where cytosine residues followed by guanine, known as CpG sites, can acquire a methyl group at the C-5 position (5mC). At the global level, a gradual demethylation of the genome is observed with age, leading to loss of genomic stability, activation of transposable elements and altered patterns of gene expression, which are themselves associated with age-related conditions. At the locus-specific level, some regions of the genome can undergo age-induced gains and losses in DNA methylation, and these changes can condition their pattern of gene expression^4^. This pattern of DNA methylation changes during the lifespan can be used as an ‘epigenetic clock’ to predict chronological age^5,6^. This phenomenon reflects deficiency in maintaining epigenetic marks and contributes to altered accessibility and aberrant gene expression which impairs the cellular and molecular functions of aged cells^5^.

Neurons, as long-lived post-mitotic cells in the brain, are characterized by an evolving epigenetic landscape during differentiation, maturation, and aging, making them a prime target for studying cellular aging in the context of brain function and health^7,8,9,10,11^. Disease signatures based on DNA methylation patterns have also been linked to a variety of age-related neurological and psychiatric disorders^12,13,14,15^. 5mC can be catalyzed to 5-hydroxymethylcytosine (5hmC) by the TET (Ten-Eleven Translocation) family of dioxygenases as part of the active DNA demethylation cycle. While much less abundant than 5mC, 5hmC levels are particularly high in the adult brain when compared to other somatic tissues^10^, accumulating during aging^16,17,18^. The precise significance of 5hmC in the brain and its accumulation during lifespan has been postulated to result from enhanced DNA demethylation activity necessary in epigenetic regulation of brain-specific genes involved in neurodevelopmental processes and in neuronal function and plasticity^19^. The dynamic changes in 5mC and 5hmC in the brain are illustrative of the different layers of complex epigenetic regulation used by neurons and other brain cells to integrate signals and outputs underlying highly skillful processes such as learning and memory.

Epigenetic mechanisms, including DNA methylation, are main actors in the regulation of monoallelic expression of gene expression, playing a crucial role in the establishment and/or maintenance of the mammalian epigenetic processes of genomic imprinting and X-chromosome inactivation^20^. Many genes regulated by these processes have critical roles in brain development and function and are thought to contribute to the diversity and specialization of neuronal cells^21^. Whether epigenetic drift affects the heritability of monoallelic expression during physiological aging and impacts the aging process remains an open question.

Imprinted genes consist of a unique subset of ∼150 genes displaying parental-of-origin-specific gene expression. The majority are located in ∼25 genomic clusters where their monoallelic expression is dependent on DNA methylation at CpG-dense regulatory regions, known as imprinting control regions (ICRs)^22^. This DNA methylation is asymmetrically deposited during female and male germline development. Interestingly, a substantial number of imprinted genes exhibit monoallelic or biased expression from one parental allele in one tissue or at specific developmental stage^23,24^. In this regard, the brain stands as one of the organs for which more genes show tissue-, isoform-or developmental-stage-specific imprinting^25,26,27^. Within the brain, this is highly regionalized with different brain areas exhibiting their own set of monoallelic or parental bias expressed genes^26,28^. The importance of imprinted genes in brain function is evidenced by the devastating neurological and behavioral conditions such as Angelman and Prader-Willi syndromes resulting from (epi)mutations affecting the chr15q11-q13 region in humans^29^. Transcriptomics studies have shown that imprinting expression in the brain is developmentally regulated, as previously appreciated for the cerebellum^25^. Whether imprinting is also susceptible to changes as a function of aging has not been systematically addressed.

X-chromosome inactivation (XCI) is a dosage compensation mechanism that equalizes X-linked gene expression of XX females to the one of XY males^30^. This process is established early in development and is regulated by the *XIST* (*X-Inactive-Specific Transcript*) long non-coding RNA (lncRNA). *XIST* is upregulated randomly from one of the two X chromosomes and becomes exclusively expressed from the inactive X chromosome (Xi). This lncRNA engages in a complex interplay with several RNA binding proteins (RBPs) to recruit transcriptional repressors and chromatin modifiers, establishing the silenced state of the Xi^31,32^. Although the molecular mechanisms underlying the initiation of X-chromosome inactivation (XCI) have been extensively elucidated, our understanding of the long-term maintenance of XCI throughout an organism’s lifespan remains limited. Recent investigations into aging, with a focus on the hematopoietic cell lineage, reveal an escalation in X-chromosome inactivation skewing, where one parental allele is preferentially inactivated^33^ along with subtle alterations in DNA methylation patterns and gene expression across the X chromosome^34,35^. Interestingly, a separate study conducted in the brain, employing single-cell transcriptomics, unveiled an intriguing finding: *Xist* expression is observed to be upregulated in aged neurons located within the hypothalamus and hippocampus of female mice^36^. The implications of this observation on X-chromosome inactivation (XCI) remain ambiguous, underscoring the necessity for in-depth investigations to elucidate the influence of aging on XCI within the context of the brain.

In the present study, we provide the first allele-specific epigenetic and transcriptional landscape of the aging mouse brain. We particularly focus on the hippocampus (HCP), a key regulator brain area of cognitive processes that tend to decline during aging. We used juvenile (8-9 weeks) and old (>100 weeks) F1 hybrid mice from reciprocal crosses between distantly related mouse strains to gain allelic resolution. DNA methylation and transcriptomic analysis enabled us to discern the impact of aging on monoallelic expression in the brain. Our results support the stable epigenetic inheritance of genomic imprinting and X-chromosome inactivation during physiological aging of the mouse brain.

## Material and Methods

### Ethics

Animal care and experimental procedures involving mice were carried out in accordance with European Directive 2010/63/EU, transposed to the Portuguese legislation through DL 113/2013. The use of animals has been approved by the Animal Ethics Committee of IMM JLA and by the Portuguese competent authority – Direção Geral de Alimentação e Veterinária – with license number 023357/19.

### Animals

Mice colonies of *Mus musculus* C56BL/6J (BL6) strain and *Mus musculus castaneus*, CAST/EiJ (CAST) strain, were obtained by the Jackson Laboratory and maintained at the Instituto de Medicina Molecular João Lobo Antunes (iMM JLA) Rodent facility. Animals were housed in a maximum of five per cage in a temperature- and humidity-controlled room (24°C, 45-65%) with a 14/12hr light/dark cycle. Animals were fed diet *ad libitum*.

Reciprocal crosses between BL6 and CAST animals - BL6/CAST (BL6 female and CAST male) and CAST/BL6 (CAST female and BL6 male) were established to generate F1 animals. Juvenile and old F1 animals were developed and sacrificed by cervical dislocation in the range of 8-9 weeks and 102-104 weeks, respectively. A total of 10 young female (6 BL6/CAST & 4 CAST/BL6), 4 young male (2 BL6/CAST & 2 CAST/BL6), 7 old female (5 BL6/CAST & 2 CAST/BL6) and 5 old male (2 BL6/CAST & 3 CAST/BL6) were used in this study (Table S1).

### Samples preparation

Sacrificed animals were decapitated by cervical dislocation, and the brains were quickly removed and the whole cerebellum (CB) and hypothalamus rapidly isolated from the brainstem. The following brain areas were then dissected according to the atlas of stereotaxic coordinates of mouse brain^37^ and immersed in liquid nitrogen for 4 seconds (s): hippocampus (HCP) (from bregma, AP: from −1.34 to −2.06 mm; ML: ±0 mm; DV: −2.8 mm); medial prefrontal cortex (from bregma, AP: from −2.10 to −1.70 mm; ML: ±0 mm; DV: −3.5 mm); and nucleus accumbens (from bregma, AP: from −1.54 to - 0.98 mm; ML: ±0.5 mm; DV: −4 mm). Lung tissue was also collected. After collection, brain areas and lung tissue were immediately frozen in liquid nitrogen and stored at - 80°C for later molecular analysis. DNA and RNA were isolated for each brain area or lung tissues from the selected animal tissues using the NucleoSpin TriPrep kit (Cat# 740966.50, Macherey-Nagel GmbH & Co.KG, Germany), following the manufacturer’s guidelines.

### 5mC/5hmC measurements by liquid chromatography-mass spectrometry

Genomic DNA from CB, HCP and lung for both juvenile and old female mice was digested using DNA Degradase Plus (Cat# E2020, Zymo Research) followed by the manufacturer’s instructions. Nucleosides were analyzed by liquid chromatography-mass spectrometry (LC-MS/MS) on a Q-Exactive mass spectrometer (Thermo Scientific) fitted with a nanoelectrospray ion-source (Proxeon). All samples and standards had a heavy isotope-labeled nucleoside mix added prior to mass spectral analysis (2′-deoxycytidine-^13^C_1_, ^15^N_2_ (Cat# SC-214045, Santa Cruz), 5-(methyl-^2^H_3_)-2′-deoxycytidine (Cat# SC-217100, Santa Cruz), 5-(hydroxymethyl)-2′-deoxycytidine-^2^H_3_ (Cat# H946632, Toronto Research Chemicals). MS2 data for 5hmC, 5mC and C were acquired with both the endogenous and corresponding heavy-labeled nucleoside parent ions simultaneously selected for fragmentation using a 5 Th isolation window with a 1.5 Th offset. Parent ions were fragmented by Higher-energy Collisional Dissociation (HCD) with a relative collision energy of 10%, and a resolution setting of 70,000 for MS2 spectra. Peak areas from extracted ion chromatograms of the relevant fragment ions, relative to their corresponding heavy isotope-labeled internal standards, were quantified against a six-point serial 2-fold dilution calibration curve, with triplicate runs for all samples and standards.

### Bisulfite treatment

Genomic DNA (1μg) from CB, HCP, hypothalamus, medial prefrontal cortex, nucleus accumbens and lung of the 4 juvenile and 4 old female mice was bisulfite converted using the EZ DNA methylation Gold kit (Cat# D5006, Zymo Research) according to manufacturer’s instructions and eluted, after column cleanup, in an elution buffer (66μl) to obtain a final concentration of ∼15ng/μl bisulfite converted DNA.

### IMPLICON library preparation and analysis

IMPLICON was performed as previously described^38^ for CB, HCP, nucleus accumbens, prefrontal cortex, hypothalamus, and lung of 4 juvenile and 4 old female mice (two animals for each reciprocal cross). Briefly, following bisulfite conversion, a first PCR amplifies each region per sample in individual reactions, adding adapter sequences, as well as 8 random nucleotides (N_8_) for subsequent data deduplication. PCR conditions and primers for this first step are listed in Table S2. Primers cover 11 imprinted clusters (10 ICRs and exon1a promoter of *Ddc* gene), together with 2 unmethylated (*Sox2, Klf4*) and 1 methylated (*Prickle1*) control regions. After pooling amplicons for each biological sample and clean-up using AMPure XP magnetic beads (Cat# A63880, Beckman Coulter), a second PCR completes a sequence-ready library with sample-barcodes for multiplexing. In this PCR reaction, barcoded Illumina adapters are attached to the pooled PCR samples ensuring that each sample pool receives a unique reverse barcoded adapter. Libraries were verified by running 1:30 dilutions on an Agilent bioanalyzer and then sequenced using the Illumina MiSeq platform to generate paired-end 250bp reads using the indexing primer with the following sequence, 5’-AAGAGCGGTTCAGCAGGAATGCCGAGACCGATCTC-3’ and 10% PhIX spike-in as the libraries are of low complexity. We run two independent IMPLICON libraries that were named: first run (lane7651) and second run (lane7950). The first run contained CB, HCP, nucleus accumbens, prefrontal cortex, hypothalamus and lung of 4 juvenile and 4 old female mice, while the second run contained samples of CB, HCP and lung of the same mice.

IMPLICON bioinformatics analysis was also performed as described^38^, following the step-by-step guide of data processing analysis in [https://github.com/FelixKrueger/IMPLICON]. Briefly, data was processed using standard Illumina base-calling pipelines. As the first step in the processing, the first 8 bp of Read 2 were removed and written into the readID of both reads as an in-line barcode, or Unique Molecular Identifier (UMI). This UMI was then later used during the deduplication step with “deduplicate bismark barcode mapped_file.bam”. Raw sequence reads were then trimmed to remove both poor-quality calls and adapters using Trim Galore v0.5.0 (www.bioinformatics.babraham.ac.uk/projects/trim_galore/, Cutadapt version 1.15, parameters:--paired). Trimmed reads were aligned to the mouse reference genome in paired-end mode. Alignments were carried out with Bismark v0.20.0. CpG methylation calls were extracted from the mapping output using the Bismark methylation extractor. Deduplication was then carried out with deduplicate_bismark, using the barcode option to take UMIs into account (see above). The data was aligned to a hybrid genome of BL6/CAST (the genome was prepared with the SNPsplit package - v0.3.4, [https://github.com/FelixKrueger/SNPsplit]). Following alignment and deduplication, reads were split allele-specifically with SNPsplit. Aligned read (.bam) files were imported into Seqmonk software v1.47 [http://www.bioinformatics.babraham.ac.uk/projects/seqmonk] for all downstream analysis. Probes were made for each CpG contained within the amplicon and quantified using the DNA methylation pipeline or total read count options. Downstream analysis was performed using Microsoft Excel spreadsheet software (v2206 Build 16. 0. 15330. 20144) and GraphPad Prism v8.0.1.

From the raw data deposited in GEO under the accession number GSE148067, the reads mapped to the following murine (mm10) genomic coordinates were excluded for consideration in this article for one of the following reasons: (1) regions that fail to reach the coverage threshold for the two parental alleles in a given sample (> 50 reads), including 3 of 13 imprinted regions, *Igf2-H19*, *Igf2r* and *Grb10,* presented in our original IMPLICON; (2) regions sequenced twice for which only the run with more reads was considered; (3) regions out of the scope of this article; for the samples sequenced in lane7651, this includes: Chr7:60005043-60005284, Chr7:142581761-142582087, Chr12:109528253-109528471, Chr6:30737609-30737809, Chr11:12025411-12025700; Chr18:12972868-12973155; for the samples sequenced in lane 7950: Chr7:142581761-142582087; Chr7:142659774-142664092; Chr11:12025411-12025700.

### RNA sequencing

Quality of Dnase I-treated total RNA from female young and old hippocampi (n= 5 old mice: n= 2 CAST-BL6 and n= 3 BL6-CAST; n= 6 young mice, n= 3 of each reciprocal cross) was checked by 2100 Agilent Bioanalyser. Samples with RIN score above 9 were processed. RNA (1 μg) was used as input for PolyA+ directional RNA-seq library preparation using the NEBNext Ultra II Directional RNA-seq Kit (#E7765, NEB) with the PolyA mRNA magnetic isolation module (#E7490, NEB) according to the manufacturer’s instructions. The pooled library was sequenced on an Illumina HiSeq 2000 with a 2×100bp kit.

Fastqs were processed using TrimGalore v0.6.6 in paired-end mode using default parameters. Validated read pairs were then aligned to the GRCm38.v5 mouse genome using Hisat2 v2.1.0 with parameters--dta--no-softclip--no-mixed--no-discordant. The resulting hits were filtered to remove mappings with MAPQ scores of < 20 and then converted to BAM format using Samtools v1.10. Allele-specific alignments were also performed by realigning mapped reads to N-masked genomes C56BL/6J (genome 1) and CAST/EiJ (genome 2) which was based on the GRCm38.v5 genome and generated using the SNPsplit v0.3.4 package. Reads that were then sorted by allele-specificity for either genome or reads containing conflicting SNP information were excluded. Total and allele specific read counts were quantified from BAMs with feature Counts (from the subread v2.0.0 package) using default parameters with gencode vM25 annotations.

We used SNP information to measure Allele-Specific Expression (ASE) and performed DESeq2 analysis between maternal and paternal alleles, excluding those genes that exhibited random (i.e., genes with monoallelic expression independent of parental origin or strain) or strain-dependent (i.e. all genes with biased expression according to the genetic background) monoallelic expression.

DESeq2 v1.34.0 package in an R (v4.1.2)/R Studio (v2022.02.0+443) environment were used thus for conducting all differential gene expression analyses (including total and allele specific analyses). For all analyses (including gene-level), lowly expressed genes were first filtered (genes with ≥10 read counts across both alleles in each sample and ≥1 TPM in each sample across all samples were kept). For allele-specific analyses, allelic ratios were first derived by dividing the maternal or paternal allele counts by the total number of allelic counts (ratios are between 0-1). These allelic ratios were then used as input into DESeq2 after adjusting sizeFactors to 1 for each sample (to account for allelic ratio input). To determine general allele specific expression (either maternal or paternal) across all mice a DESeq design (∼0 + genome + sample + allele) was used with blocking terms against the genome of origin for each allele to reduce the impact of cross-specific allelic expression. To determine allele specific expression (either maternal or paternal) in young and old mice separately, a separate DESeq2 design (∼0 + age:sample + age:allele) was utilized. Contrasts were then made between old and young mice to determine allele specific expression changes that were attributed to age. Genes with allele specific expression were considered as those with a log2FoldChange > 1 and adjusted p-value (Benjamini-Hochberg adjusted) < 0.05. For calculating the proportion of reads aligning to the mouse genome, aligned reads on ChrX were counted from BAMs using featureCounts either in an allele-specific context (post-SNPsplit for Black6 and Cast alleles) or allele-independent context (pre-SNPsplit). These counts were then divided by the total number of aligned reads and multiplied by 10^6 to obtain ChrX reads per million (RPM). ChrX count proportions for the Black6 allele were determined by dividing Black6 counts on ChrX by the total read counts on ChrX across both alleles (post-SNPsplit). Barplots, boxplots and volcano plots were plotted using the ggplot2 v3.3.5 R package, and heatmaps were constructed using the ComplexHeatmap v2.10.0 R package. The genomic distribution of imprinted genes on chromosomes were plotted using the karyoplotR v1.20.3 R package. Track plots of normalised read densities (for RNA-seq data) were plotted for several genomic loci of interest using the rtracklayer v1.54.0 and Gviz v1.38.4 R packages.

### Statistics

Statistical analysis used for each experiment is indicated in the respective figure legend with p-values indicated or marked as * p-value < 0.05, ** p < 0.01 *** p < 0.001. According to the distribution of data analyzed by the Shapiro-Wilk test, the following tests for the differential analysis of the experiments were used: unpaired two-tailed Welch’t test (Fig. 2A), two-way ANOVA followed by Tukey’s multiple comparisons test (Fig. 3B-C; Fig. S3A); Three-way ANOVA by Tukey’s multiple comparison test (Fig. S3B for the cross); and Kruskal-Wall by Dunn’s multiple comparison test (Fig. S1A-B). Differential (allelic or non-allelic) expression analysis of RNAseq data was conducted using DESeq2 package. Comparisons were considered statistically significant after a Wald test with Benjamini-Hochberg adjustment of p-values to account for multiple comparisons when adjusted p-values were < 0.05 and log2FoldChange > 1. For the GSEA analyses of differentially expressed genes, the log adj p-value for each enriched gene set and normalized enrichment score (NES) were considered for the analysis (Fig. 2Cii, Fig. S2).

## Results

### Increase in 5hmC levels is a hallmark of the aging hippocampus

To decipher the allele-specific epigenetic and transcriptional features of the aging brain, we established reciprocal crosses between BL6 and CAST mice to generate female and male BL6-CAST and CAST-BL6 F1 hybrid mice (Fig. 1). These mice were sacrificed at 8-9 weeks (young) and >100 weeks (old) of age and different brain areas as well as other organs were collected for DNA methylation and transcriptomics analyses (Fig. 1; Table S1). We concentrated our study on the hippocampus (HCP) and the cerebellum (CB) as important regions for cognitive function often impaired during aging, with lung used as a non-brain control.

**Figure 1.**
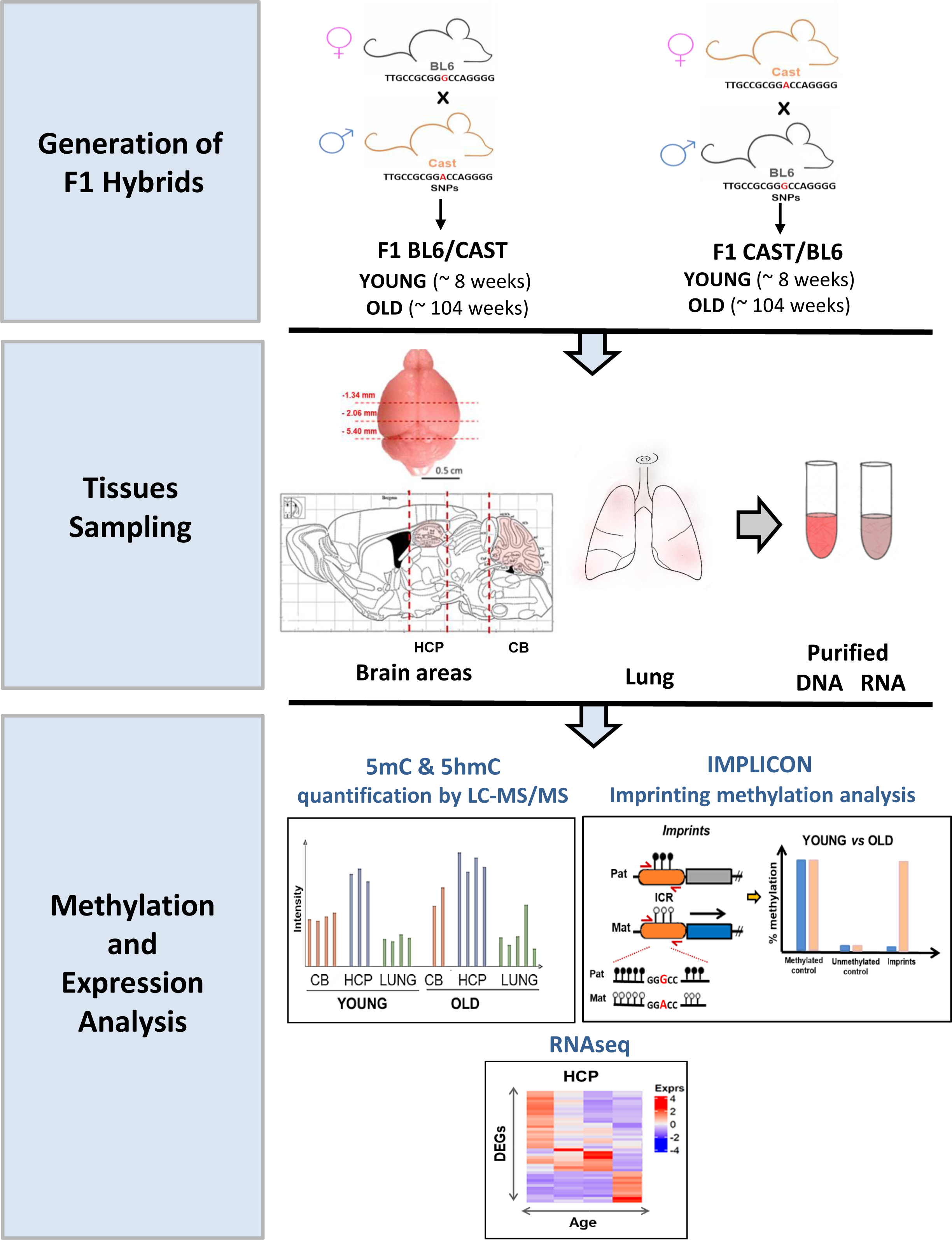
Workflow of the allele-specific methylation/transcriptional profiling of the aging brain. First, F1 hybrid mice were generated by crossing C57BL/6J (BL6) females with Cast/EiJ (CAST) males (BL6/Cast) and reciprocally crossing CAST females with BL6 males (CAST/BL6) and aged for ∼8 weeks (YOUNG) and ∼104 weeks (OLD). Second, young and old mice were sacrificed, and specific tissues were dissected such as hippocampus (HCP), cerebellum (CB) and lung. For dissection of the brain areas, stereotaxic coordinates of the mouse brain were used^37^. Dissection planes (red dotted lines) on brain tissue are shown on a sagittal view of a mouse brain from the Paxinos atlas, regions are not drawn to scale. Both DNA and RNA were extracted from the tissues and purified. Third, we perform methylation and transcriptional profiling of young and old tissue samples, by measuring global methylation levels, both 5-methylcytosine (5mC) and 5-hydroxymethylcytosine (5hmC) using Liquid Chromatography with tandem Mass Spectrometry (LC-MS/MS), by profiling allele-specific methylation at imprinting regions using IMPrint ampLICON (IMPLICON) and assessing allele-specific expression by RNA-sequencing (RNAseq).

We first evaluated the overall levels of 5-methylcytosine (5mC) and 5-hydroxymethylcytosine (5hmC) by LC-MS/MS in young and old tissues from female and male animals of both reciprocal crosses (Table S1). 5mC levels did not differ significantly among CB, HPC and lung and were not affected by age (Fig. 2Ai). In contrast, 5hmC levels were higher in the neuronal tissues, especially in HPC, where it increased further with aging (Fig. 2Aii), independently of the direction of the cross or biological sex (Fig. S1A-B and Table S3). This corroborates previous results on aging-induced increase in 5hmC levels^17,39^ and confirmed that increase in 5hmC is a hallmark of the aged HCP.

**Figure 2.**
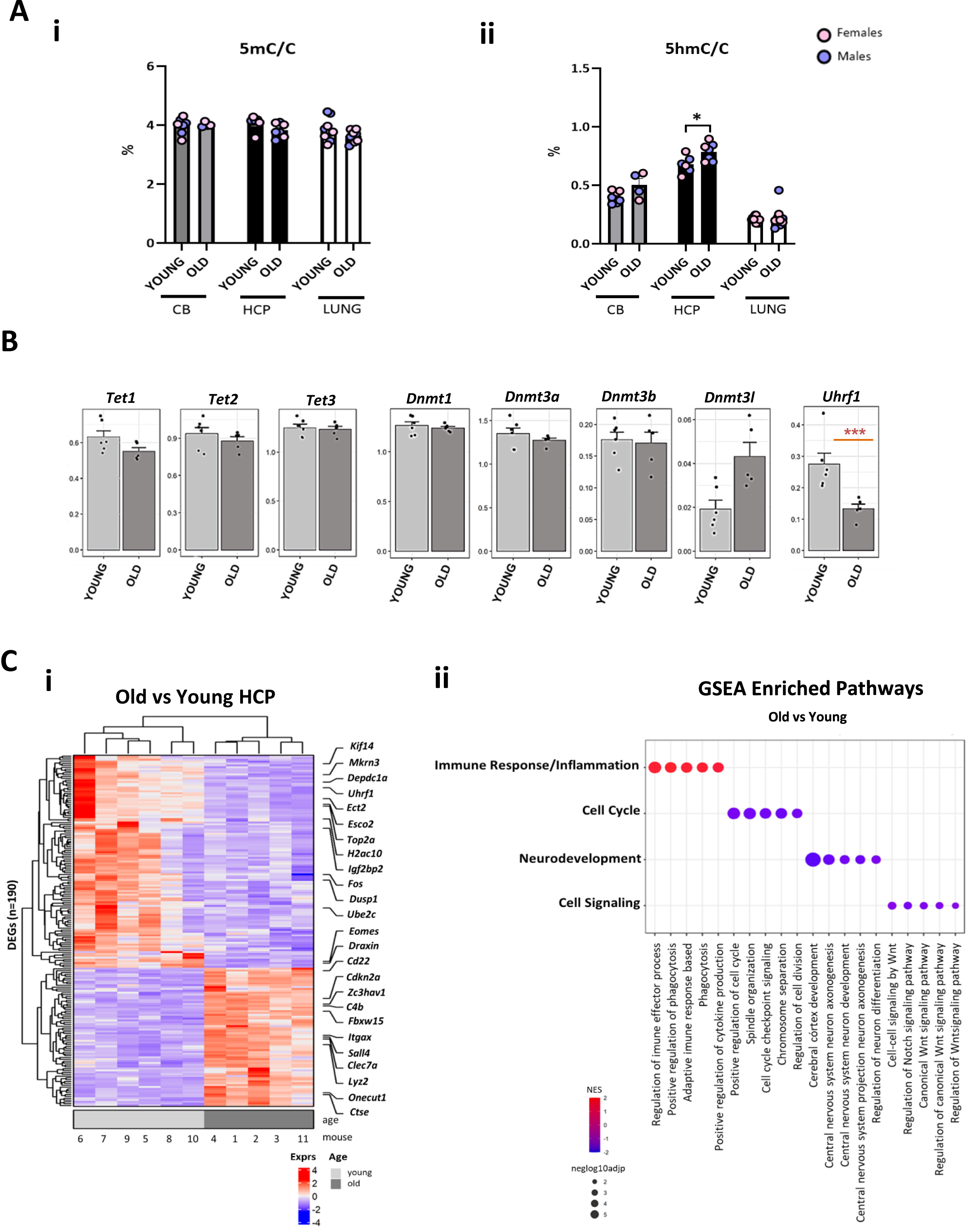
Global methylation and transcriptomics signatures of the aging brain. **(A)** Global 5-methylcytosine, 5mC, **(i)** and 5-hydroxymethylcytosine, 5hmC, **(ii)** levels measured by Liquid Chromatography with tandem mass spectrometry (LC-MS/MS) in cerebellum (CB), hippocampus (HCP) and lung of young and old female and male F1 mouse hybrids of reciprocal crosses between C57Bl/6J (BL6) and Cast/EiJ (CAST) strains. Barplots represent the average percentage of 5mC or 5hmC ± SEM of the total cytosines (C) for each the three tissues; Individual values are represented by dots and male (blue) and female (pink) mice are distinguished by colors (CB: female n=5 of which 3 young and 2 old, male n=6 of which 4 young and 2 old; HPC: female n=6 of which 3 young and 3 old, male n=7 of which 3 young and 4 old,; lung: female n=10 of which 5 young and 5 old, male n=10 of which 5 young and 5 old). Statistically significant differences are indicated as * p-value < 0.05 between young and old mice per tissue (Unpaired t-test, Welch’s correction). **(B)** Expression levels of DNA methylation/hydoxymethylation-related genes, *Tet1*, *Tet2*, *Tet3*, *Dnmt1*, *Dnmt3a*, *Dnmt3b*, *Uhrf1* measured by RNAseq. Barplots represent the average log10(TPM+1) expression levels ± SEM of the different genes in young and old female HCP (n=6 young mice; n=5 old mice). Statistically significant differences determined by DESeq2 (Wald tests with BH-adjustment of p-values to account for multiple comparisons) are indicated as *** adjusted p-value < 0.05 and |log2 (fold-change) | >1. **(C)** Transcriptional phenotypes of the aging hippocampus in mice Heatmap plot of the differentially expressed genes (DEGs) of young versus old hippocampi. Horizontal axis represents each mouse number ordered by age (light gray - young - total of 6 female mice; dark gray - old - total of 5 female mice); DEGs (n = 190 genes) determined by DESeq2 (Wald tests with BH-adjustment of p-values) were selected by adjusted *p-value* <0.05 and |log2 (fold-change) | >1. Gradient of color from red to blue, denotes, respectively, upregulated, and downregulated DEGs in young vs old HCP. Examples of a few genes that are UP or DOWN are shown on the left side of the heatmap, **(i)**; Dot plot showing representative GSEA pathways that were highly enriched within ontology gene sets using DEGs determined above (M5 pathway collection from the mouse Molecular Signatures Database (MSigDB)). Pathways are grouped into four groups where dot size represents the-log adj p-value for the enriched gene set. Normalized Enrichment Score (NES) > 0 (red) indicate pathways enriched in old mice, and NES < 0 indicate depleted pathways in old mice (blue), **(ii)**.

### Transcriptomic signatures of the aged hippocampus

We next examined alterations in the transcriptome occurring during the aging of the HCP by employing RNAseq on samples obtained from both young and elderly subjects. We selected HCP due to the rise in 5hmC levels associated with aging. Specifically, we performed RNAseq analyses on female HCP, comprising a total of 5 samples from aged mice (2 CAST-BL6 and 3 BL6-CAST individuals) and 6 samples from young mice (with 3 individuals from each reciprocal cross). This dataset was thoroughly scrutinized to assess shifts in the expression of genes associated with the DNA methylation machinery over the aging process, as well as to investigate broader patterns of gene expression changes.

To better understand the cause leading to the increase of 5hmC levels upon aging, we first analyzed expression levels of the ten-eleven translocation (Tet) methylcytosine dioxygenases (*Tet1*, *Tet2* and *Tet3*) responsible for the sequential conversion of 5mC to 5hmC, and subsequent oxidation steps as part of the active DNA demethylation pathway^40^. No differences were observed for any of the three *Tet* genes by RNAseq in HCP (Fig. 2Bi). Likewise, no differences in the DNA methyltransferase genes (*Dnmt1*, *Dnmt3a*, *Dnmt3b* and the *Dnmt3l* cofactor), responsible for catalyzing the transfer of a methyl group to DNA, were observed between young and old HPC (Fig. 2Bi). Curiously, a significant drop in *Uhrf1* mRNA expression, a gene encoding for a DNMT1-interacting protein essential for maintenance of methylation through DNA replication^41^, was observed in the aged HPC (Fig. 2B). *Uhrf1* downregulation was not associated with measurable changes in 5mC levels perhaps because aged HPC is composed mostly by postmitotic cells. Overall, we rule out that the core DNA methyl/hydroxymethyl machinery is affected by aging.

Next, to conduct a broader analysis of gene expression, we performed a differential gene expression analysis, which identified 75 upregulated and 115 downregulated genes during the aging process (adjusted *P* value (*P*adj) < 0.05, fold change (FC) > 1) (Fig. 2Ci; Table S4). Within the set of upregulated genes, we observed the presence of genes associated with cell cycle inhibition, including *Cdkn2a*, as well as several genes involved in immune response and inflammation, such as *Cd22*, *Clec7a*, *Ctse*, *Lyz2*, and *C4b*. Accordingly, previous single-cell RNAseq analyses have identified *Lyz2* as an aging marker of microglia and *C4b* as an aging marker of astrocytes^36^. Downregulated genes were involved in cell cycle progression and chromosome segregation (e.g., *Kif4*, *Ect2*, *Esco2* and *H2ac10*), regulation of transcription (e.g., *Fos* and *Top2a*) and neural development (e.g., *Eomes* and *Draxin*). To complement this analysis, we performed Gene Set Enrichment Analysis (GSEA) to capture changes in the expression of pre-defined sets of genes and investigate further the cellular processes altered with age (Table S5). GSEA identified 4 major cellular processes that were altered upon age. Three cellular processes consistently exhibited downregulation: cell cycle, cell signaling, and neurodevelopment. Conversely, immune response and inflammation were consistently upregulated (Fig. 2Cii). Within these four altered cellular processes, the top five gene sets for each major category are depicted in Fig. 2Cii and comprehend “Positive regulation of cell cycle”, “Cerebral cortex development”, “Cell-cell signaling by *Wnt*” as downregulated pathways and “Regulation of immune effector process” as an upregulated pathway (Fig. 2Cii; Fig. S2). Overall, our transcriptomics analysis confirmed an aging transcriptional signature in the aged HCP samples.

### Imprinting methylation is stable during aging

Next, we investigated how aging influences the fidelity of genomic imprinting. We first screened for imprinting methylation using the IMPLICON method that we previously developed and validated on tissue samples of F1 mice from reciprocal BL6xCAST crosses^38^. IMPLICON is an amplicon-sequencing method measuring DNA methylation at several ICRs across the genome at the nucleotide resolution with high coverage^38^. With this method, we are also able to separate out paternal from maternal reads based on SNPs between BL6 and CAST strains that are contained in our amplicons and conserved during bisulfite conversion. IMPLICON was successfully performed on 11 imprinted clusters (10 ICRs and the exon1a promoter of *Ddc* gene) together with 2 unmethylated and 1 methylated control regions (Fig. 3A, Table S6) in HCP, CB and lung of four young and old female mice from both F1 BL6xCAST reciprocal crosses.

**Figure 3.**
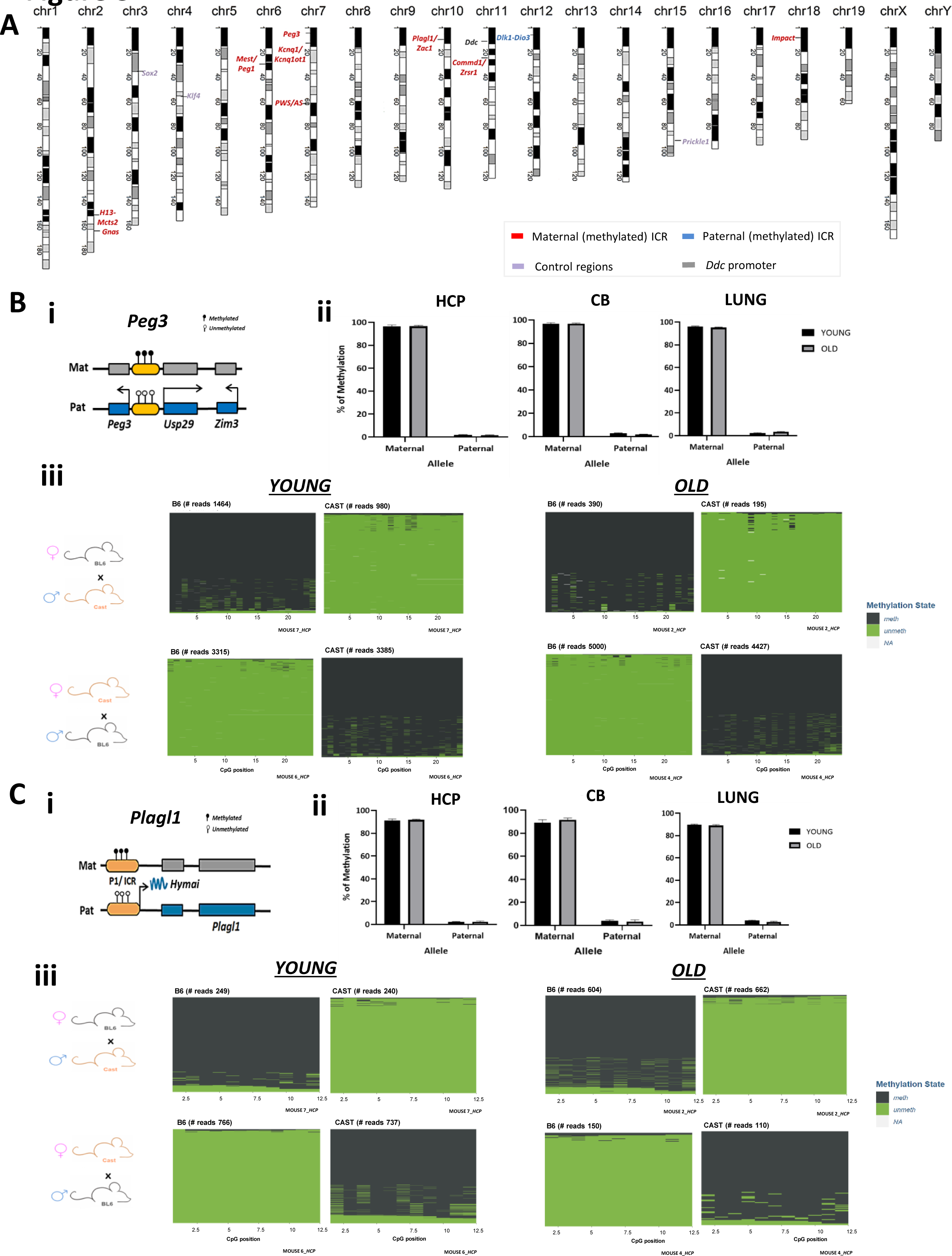
IMPLICON reveals methylation stability at imprinted loci. **(A)** Schematic view of the murine karyotype depicting the location of the regions studied by IMPLICON; red and blue letters mark imprinted regions with maternally inherited and paternally inherited methylation respectively; violet and gray letters mark control regions and Dopa decarboxylase (*Ddc*) promoter, respectively. **(B-C),** IMPLICON analysis of *Peg3* **(B)** and *Plagl1* **(C)** imprinted clusters in hippocampus (HCP), cerebellum (CB) and lung of 4 juvenile and 4 old F1 female mouse hybrids of C57Bl/6J (BL6) and Cast/EiJ (CAST) reciprocal crosses (2 animals of each reciprocal cross per age group) measured by IMPLICON; Scheme on the left of each graph represents the expected methylation status of each region (white lollipops – unmethylated CpGs; black lollipops – methylated CpGs; gray box - non-expressed gene; blue box - paternally expressed gene; Mat – maternal allele; Pat – paternal allele; regions are not drawn to scale. For *Plagl1*, in the scheme *Hymai* noncoding RNA is also represented, being an exon gene with the transcription starting site from the same ICR promoter **(i)**. Barplots represent the mean ± SEM methylation levels measured at each CpG within different genomic regions per parental allele in F1 female mice for CB, HCP, and lung; **(ii)**. Descriptive plots displaying methylated (green) and unmethylated CpGs (gray) for each CpG position (in columns) in all the individual reads (in rows) for both the ICR of both *Peg3* and *Plagl1* imprinted loci in the HCP of 2 young and 2 old female mice from each reciprocal cross; NA-means not applicable, when methylation status was not retrieved **(iii)**.

Unmethylated (*Klf4 and Sox2)* and methylated (*Prickle1*) controls, showed low (<∼10%) or high (>∼90%) DNA methylation levels respectively at both maternal and paternal alleles for all tissues analyzed (Fig. S3A; Table S6) irrespective of age. We next investigated DNA methylation levels of ICRs of imprinted regions. Our results confirmed that DNA methylation at all ICRs analyzed was stably maintained with age in the HCP, CB and lung (Table S6). This is illustrated for the maternally methylated *Peg3* and *Plagl1* loci (Fig. 3Bii-Cii), which showed >90% DNA methylation of the maternal allele and <10% DNA methylation of the paternal allele, irrespective of the genetic background (Fig. S3B).

We are also able to examine DNA methylation consistency along individual reads using the IMPLICON method. We confirmed reads were either fully unmethylated or methylated depending on their parent of origin, independently of the genetic background. This is exemplified for *Peg3* and *Plagl1* ICRs in HCP (Fig. 3Biii-Ciii). Taking advantage of the single nucleotide resolution of IMPLICON, we also determined methylation levels at each CpG within each genomic region with aging but did not observe consistent and meaningful differences in the various tissues across aging (Fig. S3C; Table S6).

Using IMPLICON, we also checked the methylation levels of the *Dopa decarboxylase* (*Ddc)* gene, which is involved in dopamine biosynthesis, and often dysregulated in neurodegenerative and psychiatric disorders^42^. *Ddc* transcripts are generally expressed from both parental alleles in most of the body tissues but can also exhibit isoform-, cell-, development-specific imprinted expression including in the brain^43,44^. Here we reported that the exon1a promoter of the imprinted gene isoform of *Ddc* was not differentially methylated between the parental alleles, nor throughout aging, in HCP and CB, with methylation levels in the two brain areas being lower than in the lung (∼50% vs 90%) (Fig. S3Aiii).

Finally, we also extended our IMPLICON analysis to three additional brain regions. Similar to CB and HCP, the nucleus accumbens, prefrontal cortex and hypothalamus showed high fidelity of differential DNA methylation for the 6 imprinting clusters and the control regions analyzed (Table S7). In conclusion, our results suggest that imprinting methylation is stably maintained during aging for the loci and brain areas investigated in this study.

### The “*Imprintome*” of the young and old hippocampus

We next determined the “*imprintome*” of the aging mouse HPC by taking advantage of our transcriptome dataset from F1 mice of reciprocal crosses. In total, we identified 113 genes with parental-of-origin monoallelic expression in young HCP, of which 66 were maternally and 49 were paternally expressed. These genes were in 17 genomic regions that corresponded to known imprinted regions (Fig. 4A-B; Table S8).

**Figure 4.**
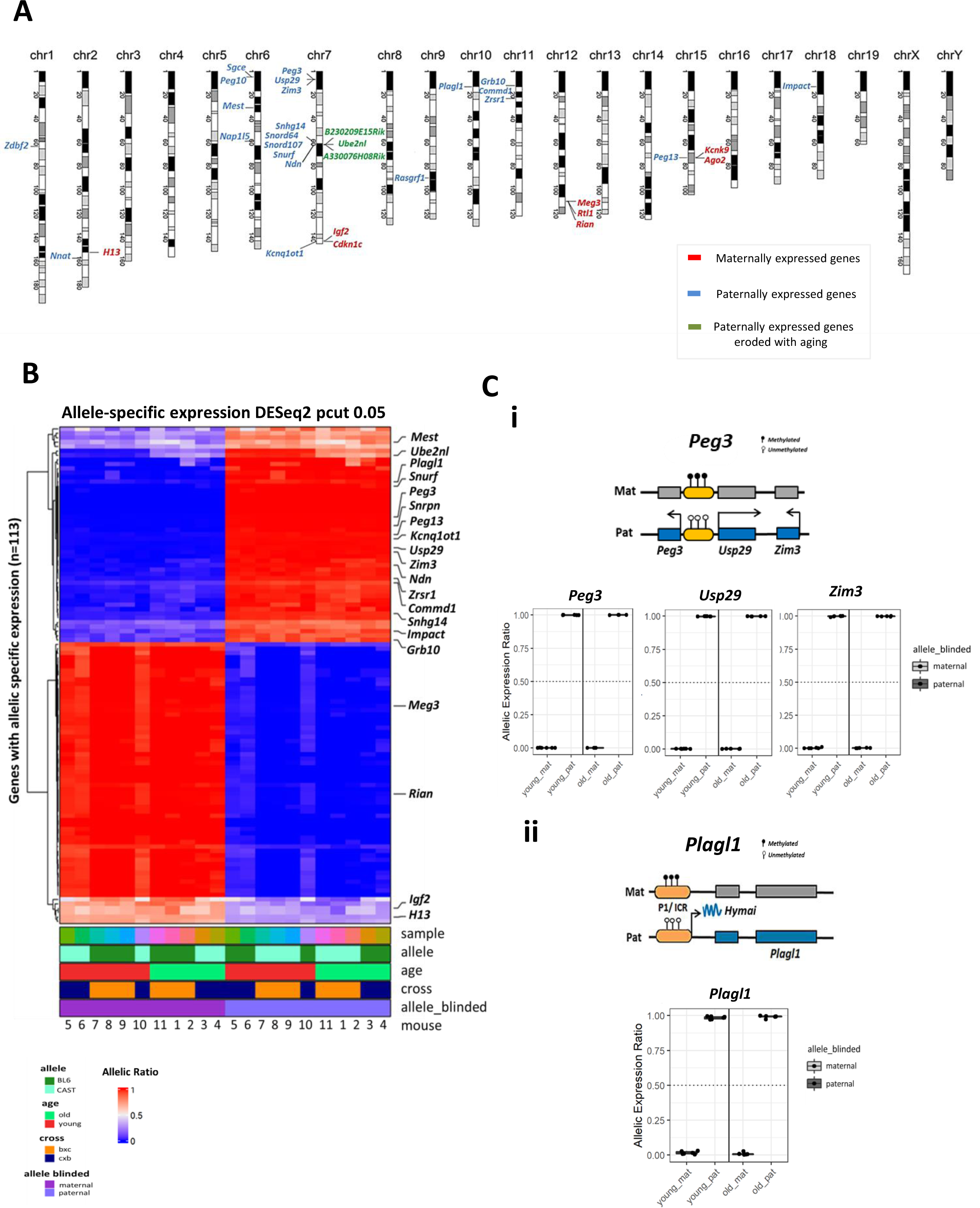
The “*imprintome* “of the murine HCP. **(A)** Schematic view of the murine karyotype depicting the location of the imprinted genes detected by RNAseq in HCP; maternally expressed imprinted genes are marked in red; while and paternally expressed imprinted genes are marked in blue; genes marked in green are paternally expressed genes that erode with aging; for sake of simplicity small RNAs, such as microRNAs and snoRNAs, pseudogenes and noncoding genes of unknown function are not represented, with the exception of the genes marked in green; the full list of imprinted genes detected by RNAseq in HCP is in Table S8. **(B)** Heat map depicting the results of of RNAseq transcriptomic analysis illustrating for differential parental-specific gene expression of HCP in young (n=6) and old (n=5) female mice (F1, n=3 young and n=3 old mice; and reciprocal cross, n=3 young and n=2 old mice). Color corresponds to per-gene allelic expression scores (maternal counts/total allelic counts) which are between 0-1, scores > 0.5 (red) or < 0.5 (blue) represent genes with a maternal or paternal expression bias, whereas those with ∼0.5 represents genes with bi-allelic expression. Only genes were hierarchically clustered based on these allelic ratios using euclidean-distance and ward. D2 based clustering. The scales on the bottom show the color codes for allelic expression levels. Examples of genes with maternal or paternal allelic expression biases are annotated on the right side of the heatmap. **(C)** Boxplot representing median allele-specific expression of the parental allele of the *Peg3, Usp29, Zim3,* **(i)** and *Plagl1,* **(ii)** imprinted genes in the HCP of young (n=6) and old (n=5) female mice measured by RNAseq (DESeq2, *P*adj < 0.05, log2FC > 1). Boxplot boundaries indicate the lower (1^st^) and upper (3^rd^) quartiles of allele expression ratios, with the middle line indicating the median value. Scheme on top of each graph represents the expected methylation status of *Peg3* and *Plagl1* imprinted regions (white lollipops – unmethylated CpGs; black lollipops – methylated CpGs; gray box - non-expressed gene; blue box - paternally expressed gene; Mat – maternal allele; Pat – paternal allele; regions are not drawn to scale. For the *Plagl1* locus, in the scheme *Hymai* non-coding is also represented, being an exon gene with the transcription starting site from the same ICR promoter. Drawn not in scale.

We next investigated whether the “*imprintome*” of the mouse HCP changed upon aging. Consistent with our IMPLICON results, RNAseq analysis revealed stable imprinting expression during aging in HCP. For example, *Peg3, Usp29, Zim3* and *Plagl1* genes are exclusively paternally expressed in young and old HCP (Fig. 4Ci-ii; Fig. S4A-B), while *Ddc*, which shows no differential methylation between the parental alleles, is expressed from both alleles in both young and old HPC (Fig. S4C). In contrast, its neighboring imprinted gene, *Grb10* was consistently expressed from the paternal allele as expected not only in young but also in old HCP. In line with the lack of aging-specific effects on genomic imprinting, expression levels of genes implicated in imprinting maintenance such as *Dppa3*, *Trim28*, *Zfp445* or *Zfp57* remained constant. Although there was a tendency for decreased *Zfp57* expression in old HCP, this trend was not statistically significant (Fig. S4D).

More globally, when comparing old versus young ASE for imprinted genes using DESeq2 method, we found three not previously reported imprinted genes (*B230209E15Rik*, *Ube2nl* and *A330076H08Rik*) showing aging-specific partial erosion of strict pattern of monoallelic expression (deviating from a maternal:paternal ratio of ∼0: ∼100% in young to ∼25:∼75% in old HPC) (Fig. 5A; Table S9). Both *B230209E15Rik* and *A330076H08Rik* encode for noncoding transcripts, while the pseudogene *Ube2nl* (Ubiquitin Conjugating Enzyme E2 N Like) has 91% conservation with the gene encoding for the multi-exonic ubiquitin-conjugating enzyme *E2N*. Their loss of strict monoallelic expression in old HCP had a minor impact in the overall expression level of these genes (Fig. 5B) but was a consistent feature of the aged HCP. All these genes are located within the PWS/AS imprinted cluster on chr7, with the *Ube2nl* pseudogene situated within an intron of *B230209E15Rik* (Fig. 5C). They are normally expressed from the paternal allele and are located between the paternally expressed *Snurf-Snrpn* gene and the proximal side of the cluster that includes *Mkrn3*, *Magel2* and *Ndn* genes. Interestingly, it is in the PWS/AS locus, where we find the only imprinted gene, *Mkrn3* which is downregulated in old HPC, but, in this case, does not alter its RNA allele-expression ratio (Fig. S5A). In conclusion, these transcriptomics data is consistent with the IMPLICON results and reveals remarkable stability of genomic imprinting in physiological aging, with the rare exception of three transcripts within the large PWS/AS imprinted cluster.

**Figure 5.**
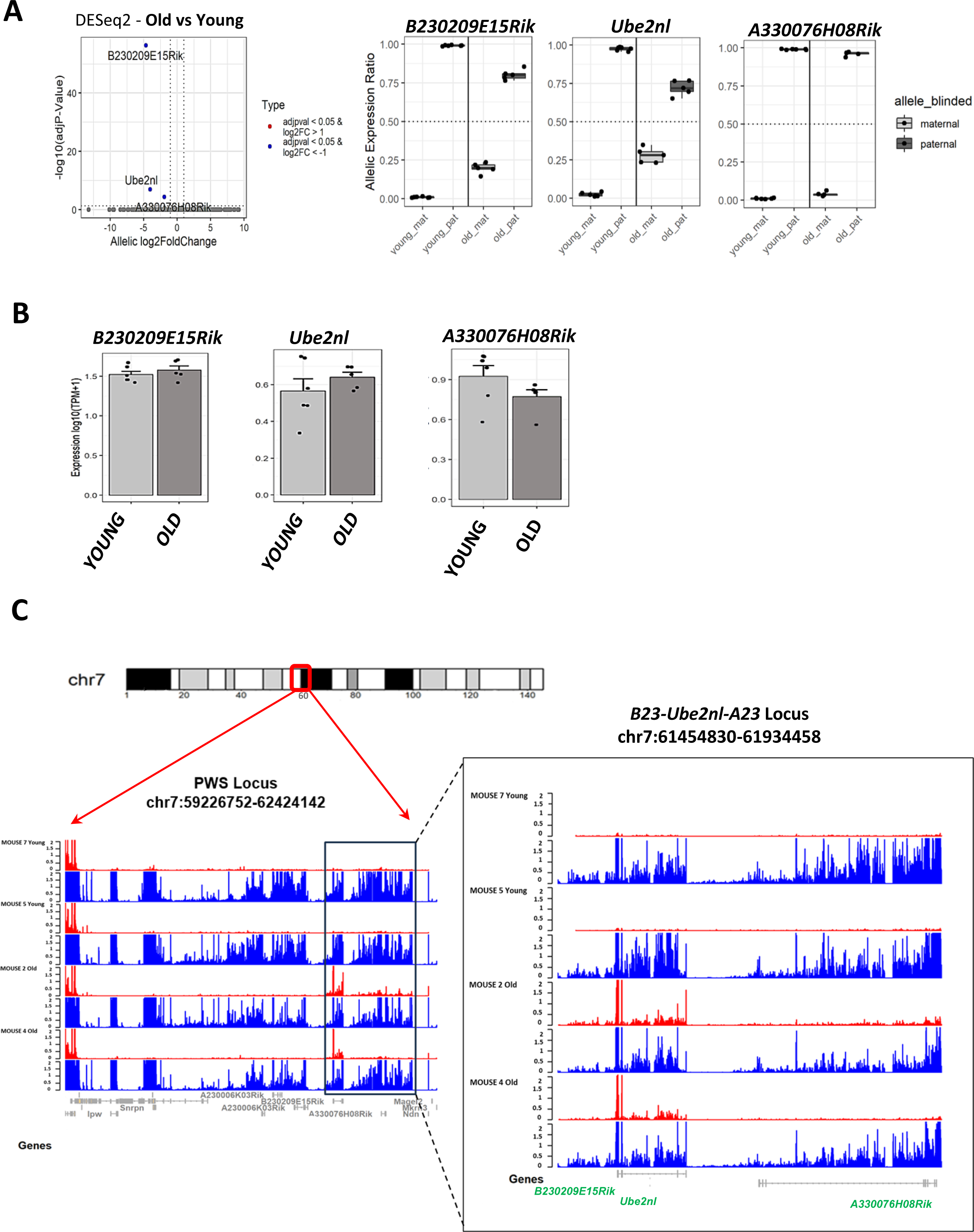
Loss of strict monoallelic expression of three imprinted noncoding genes within the Prader-Willi syndrome/Angelman syndrome locus. **(A)** On the left, dotplot detecting the allelic expression changes between young and old HPC on the newly reported imprinted genes (*B230209E15Rik*, *Ube2nl* and *A330076H08Rik*) located Prader-Willi syndrome/Angelman syndrome (PWS/AS) locus using DESeq2 (Wald test and BH-adjustment of p-values) method comparing old vs young (*P*adj < 0.05, |log2FC| > 1). The log2FC represents the change in allele expression ratios between old and young mice for each gene. log2FC< -1 represents those genes which lose allele-specific expression, whereas log2FC> 1 represents those genes which have gained allele specific expression; on the right, boxplots represent the median allelic expression ratio of the parental alleles for the genes *B230209E15Rik*, *Ube2nl* and *A330076H08Rik* in young (n=6) and old (n=5) female HCP. Boxplot boundaries indicate the lower (1^st^) and upper (3^rd^) quartiles of allele expression ratios, with the middle line indicating the median value. **(B)** Bulk expression levels of *B230209E15Rik*, *Ube2nl* and *A330076H08Rik* of young and old hippocampi. **(C)** Representative image of the PWS/AS locus in the mouse chr7 and RNA-seq allelic tracks representation (red = maternal, blue = paternal), 2 young and 2 old mice were chosen for display (each mouse from reciprocal cross), showing the location of the three novel imprinted genes differentially modulated in aged HCP Region in square indicates the *Ube2nl/B230209E15Rik (B23)/ A330076H08Rik (A23)* locus within the PWS/AS region, where expression from the maternal allele becomes evident in the older mice.

### X-chromosome inactivation in the aging brain

XCI is a dosage compensation mechanism that silences one of the two X chromosomes in female mammalian cells^31^. We utilized our RNAseq dataset, which includes young and old female F1 hybrid mice from reciprocal crosses, to examine the status of X-chromosome inactivation (XCI) during the aging process in HCP. First, we assessed the expression levels of the long noncoding RNA *Xist* (*X-inactive-specific-transcript*) which is the master-regulator of XCI. There were no differences in *Xist* levels between young and old HCP (Fig. 6A). Likewise, the expression of other noncoding genes located on the X-inactivation center, such as *Ftx*, *Jpx*, and *Tsix*, remained unchanged with aging, which contrasts with the findings of a recent single-nuclei RNAseq study^36^ (Fig. S6A). Next, we explored the allele-specific expression of *Xist* and genes on the X chromosome using BL6/CAST SNP information, to gain insights on potential effects of aging on XCI. We observed a bias towards the expression of the BL6 *Xist* allele irrespective of parental origin or age, with some variation between individual animals (Fig. 6Bi-ii; Fig. S6B). Increase in *Xist* expression from the BL6 allele resulted in a concomitant reduction of expression from across the entire BL6 X chromosome (Fig. 6Bi-ii). This suggested preferential inactivation of the BL6 X chromosome which is consistent with previous findings reporting a skew towards the inactivation of the BL6 X chromosome in BL6-CAST hybrid female mice^45^. When analyzing individual genes, we observed a decrease of the percentage of BL6 reads for X-linked genes such as *Mecp2*, *Chic1*, *Atrx* or *Huwe1* (Fig. 6C; Fig.S6C). This was a generalized behavior for all X-linked genes which include escape genes such as *Eif2s3x* and *Kdm5c* (Fig. S6D). Although they can be expressed from both active and inactive X chromosomes (Fig. S6D), this pattern suggests that escape genes are generally more prominently expressed from the active X chromosome than from the inactive one. Some genes evade this general allelic trend across the X chromosome. For example, *Pgk1* gene, traditionally used as a biomarker of X-inactivation status^46,47^, exhibits a strong preference for the expression of the CAST allele. In contrast, *Firre*, a non-coding RNA with an important role in conformational organization of the Xi^48,49^, exhibited strong BL6 biased expression (Fig. S6D). Finally, as a proxy for relaxation of XCI, we measured whether there was any increase in gene activity from the X chromosome. There was a marginal but not statistically significant increase (*Padj* = 0.51) in the proportion of reads mapping to the X chromosome in aged HCP, consistent with what we observed for individual X-linked genes (Fig. 6D; Fig S6A). Overall, our results suggest X-chromosome inactivation is stable, with no signs of exacerbated skewing nor Xi relaxation, during physiological aging of the HCP.

**Figure 6.**
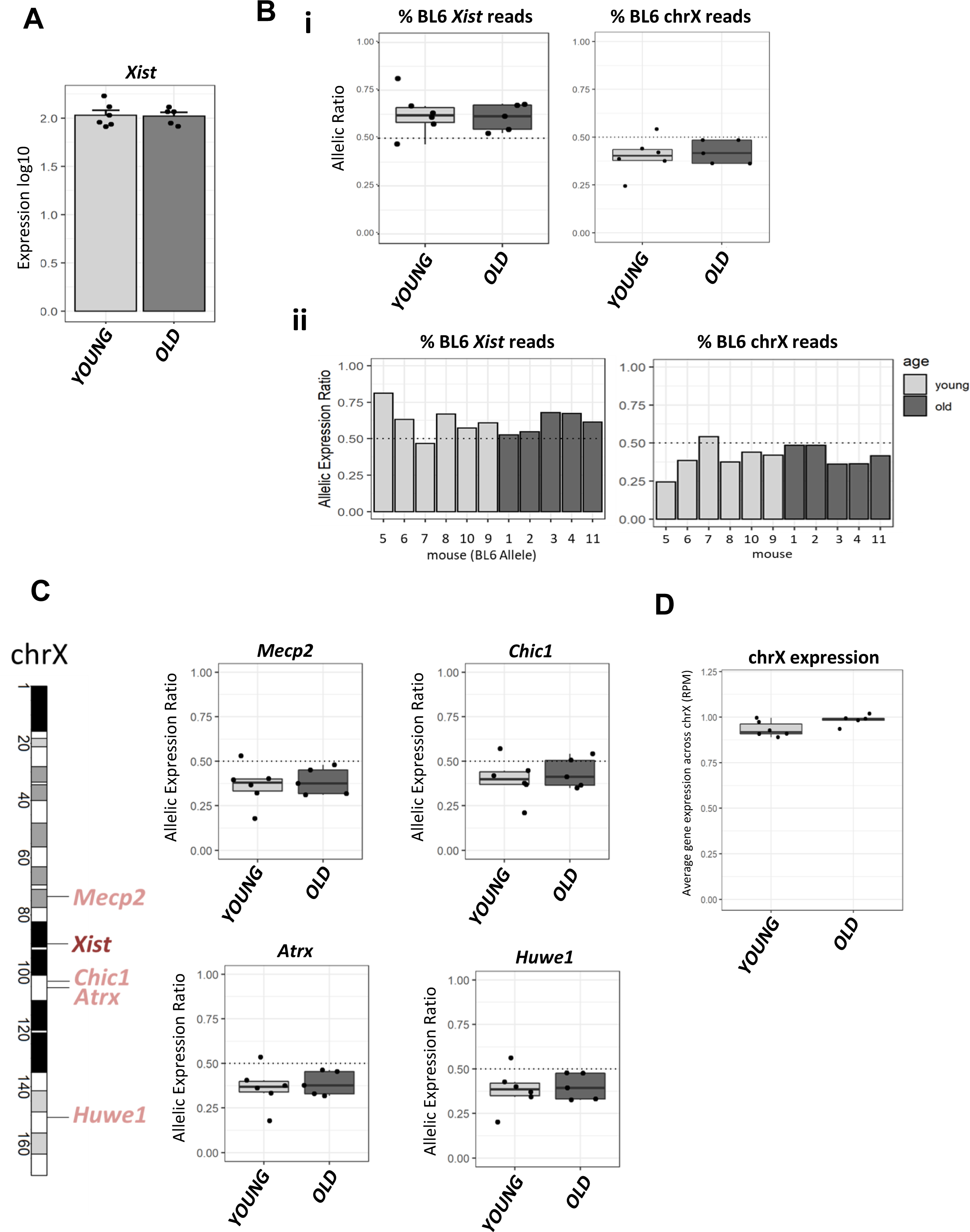
Stability of X-chromosome inactivation in the aged HCP. **(A)** Expression levels of *Xist* (X-inactive-specific-transcript) long noncoding RNA in young (n=6) and old (n=5) female mice in log10(TPM+1) (transcripts per million) measured by RNAseq. **(B)** Allele-specific expression ratio of *Xist* gene and of the entire chrX reported as percentage of BL6 reads averaged for young (n=6, of which 3 BL6/CAST and 3 CAST/BL6) and old mice (n=5, of which 3 BL6/CAST and 2 CAST/BL6)) in boxplots **(i)**, and for each individual mouse **(ii)**. Boxplot boundaries indicate the lower (1^st^) and upper (3^rd^) quartiles of allele expression ratios, with the middle line indicating the median value. **(C)** Median allelic expression ratio of X-linked genes; *Mecp2*, *Chic1*, *Atrx* or *Huwe1* measured by RNAseq represented by barplots depicted the % of BL6 reads. Boxplot boundaries indicate the lower (1^st^) and upper (3^rd^) quartiles of allele expression ratios, with the middle line indicating the median value. On the left, scheme of the chrX with the location of *XIST* and these genes. **(D)** Boxplot graph representing the median gene expression across chrX in reads per million (RPM) for young and old mice. Boxplot boundaries indicate the lower (1^st^) and upper (3^rd^) quartiles of allele expression ratios, with the middle line indicating the median value.

## Discussion

In this study we present the first allele-specific epigenetic and transcriptional landscape of the aging mouse brain, with a specific emphasis on the HCP. Using a combination of IMPLICON and RNA sequencing analyses, our findings highlight the robust preservation of genomic imprinting and the stability of X-chromosome inactivation throughout the natural aging process in the murine brain.

Epigenetic drift, a key hallmark of aging, has been implicated in the irreversible decline of organismal fitness and the onset of aging-related illnesses^50^. Notably, epigenetic modifications, including DNA methylation, undergo predictable changes over time, enabling the estimation of biological age based on DNA methylation ‘*epigenetic clocks*’^51^. Whether these epigenetic-induced changes affect maintenance of monoallelic expression including genomic imprinting and X-chromosome inactivation, was not known and thus was the focus of this article.

One feature of epigenetic drift in the brain is the increase in 5-hydroxymethylation (5hmC) levels over time^52^. Indeed, 5hmC levels increase markedly during lifespan, suggesting that 5hmC-mediated epigenetic modification may be critical in neurodevelopment and neurodegenerative disorders^53,54^. Our data confirmed that 5hmC levels are higher in the brain than in peripheral tissues and increases in the HCP with age, irrespective of biological sex^55,53^ (Fig. 2A; Fig. S1A). Previous studies have similarly reported age-associated increases in 5hmC in CB^57,58,59^. In our study, we report a trend towards increased 5hmC levels in CB, but it was not statistically significant, potentially due to the small number of CB samples analyzed (Fig. 2A; Table S1). In the HCP, the observed increased 5hmC signal during aging occurred in the absence of any changes in 5mC levels, suggesting that 5hmC may be acting as an epigenetic marker rather than an intermediary in DNA demethylation^19^. Why 5hmC levels increase during aging in the brain is not yet fully understood. One hypothesis may be that 5hmC levels increase as a compensatory mechanism in response to age-induced changes due to cellular stress in the brain’s microenvironment^60,61^. Increased 5hmC levels are generally associated with enhanced activity of Tet dioxygenases^59^, however, expression level of these enzymes in HCP did not change over time, consistent with previous studies^62^. This suggests that the increase in 5hmC may not rely on expression levels but perhaps driven by altered enzymatic activity^61^. One interesting observation from our study was a decrease of *Uhrf1* transcript levels during aging. While Uhrf1 is primarily associated with the recognition and maintenance of 5mC during DNA replication, it can also interact with 5hmC and influence the recruitment of DNA methyltransferases^63,41^. Interestingly, previous research has demonstrated that loss of Uhrf1 lead to a global increase in 5hmC levels^41^. However, the specific mechanism underlying this differential regulation between Uhrf1 and 5hmC remains unclear and requires further investigation.

Our transcriptome analysis identified both upregulated and downregulated transcripts in the aged HCP. Notably, we found higher expression of genes related to inflammation and immune responses, while genes involved in cellular cycle progression, neurodevelopmental processes and active signaling pathways displayed reduced expression levels (Fig. 2Cii, Fig. S2B). As previously reported^64,51^, the upregulation of inflammatory pathways^62^ and downregulation of genes involved in pathways of growth factor signaling, encoding mitochondrial proteins and protein synthesis machinery^51^ are common characteristics of aging in the brain. Interestingly, we observed canonical Wnt (Wnt/β-catenin) signaling to be dysregulated with aging. Increasing evidence indicates that Wnt signaling regulates multiple aspects of adult hippocampal neurogenesis^65^ as well as neural function and synaptic connectivity^66^ and downregulation of Wnt signaling could be involved in the cognitive decline associated with aging and Alzheimer’s disease^67^. Together, our transcriptome analysis aligns with previous studies^68,36^, suggesting a consistent and generalized pattern of gene expression changes associated with aging.

Genomic imprinting is an enduring form of epigenetic inheritance established in parental germ cells and maintained throughout an organism’s development^24^. In the central nervous system, imprinting is important for neurogenesis, brain function and behavior^27^ and dysregulation of imprinting results in neurodevelopmental and behavioral disorders such as PWS/AS^29,24^. In accordance, the brain, especially neurons, consistently shows a high number of imprinted genes in adulthood^18,26,69^. A detailed investigation of imprinted gene expression in the aging HCP was yet to be performed, despite previous association between imprinting methylation and hippocampal volume in aging^70^. Recent evidence suggested that imprints can be dysregulated by environmental insults during critical periods of embryonic/fetal development^71,72,73^ with long-term consequences in tissue function and susceptibility to age-related diseases^25,26^. Whether imprinting in the brain is also susceptible to changes as a function of aging has not been addressed previously in a systematic way.

Using our allele-specific IMPLICON method, we observed a consistent DNA methylation pattern across 11 imprinted regions in the HCP, CB, and lung tissues of both young and old F1 mice, as well as in their reciprocal crosses. We also noted, however, that certain CpGs exhibited either increased or decreased methylation. Whether this is attributed to experimental noise or indeed has biological meaning would require further investigation. We also assessed age-related allele-specific transcriptional changes in the HCP in our RNAseq dataset, documenting 113 coding and non-coding genes with parental allelic expression in the HCP across aging. In line with our IMPLICON findings, imprinted gene expression remained stable throughout the aging process. While we cannot rule out dysregulation of imprinting in the context of age-onset diseases or following environmental insults^74,75^, our findings show stable parental-of-origin DNA methylation and transcription at imprinted loci during physiological aging of the brain. One caveat in our analysis is the absence of single-cell resolution which masks the existence of other imprinted genes which may exhibit cell type-specific imprinting. This may indeed be the case of *Ddc* gene belonging to the *Grb10* imprinted locus on chr11, which we postulated to undergo transcriptional or imprinting regulation upon aging and thus influencing dopamine production^76,43,44^. Single-cell techniques, such as scRNAseq, would be required to disclose the full “*imprintome*” of HCP during aging. However, scRNAseq still holds limitations for an accurate allele-specific map of the transcription due to insufficient read depth and frequent allele dropouts which need to be improved in the future.

Although our data strongly points for an enduring stability of genomic imprinting in the aging brain, an exception was found for three novel non-coding transcripts at the PWS/AS imprinted locus on mouse chr7 that lost strict monoallelic expression upon aging: *B230209E15Rik*, *Ube2nl*, *A330076H08Rik*. These transcripts are strongly expressed in brain tissues; however, their exact function is currently unknown. The most enigmatic of them all is *Ube2nl*, an intronless pseudogene with an open reading frame derived from the *Ube2n* (ubiquitin-conjugating enzyme E2N) gene on mouse chr10. While we observed consistent allele biases changes in the aged HCP for these genes, this did not translate into overall gene expression changes. Functional studies will be needed to understand the role of these transcripts in mouse development and aging and whether the syntenic region in humans is also sensitive to aging-specific effect on imprinting regulation.

We also investigated changes in X-chromosome inactivation (XCI) during the aging process, and unlike recent single-cell RNA sequencing results indicating unexpected *Xist* upregulation in the hypothalamus and to some extent in the HCP^36^, we did not observe any differences in our cohort of 11 animals (Fig. 6A). This also extended to other genes on the X-inactivation center, *Ftx*, *Jpx*, and *Tsix* which also did not change with aging (Fig. S6A). This disparity might be explained by the differences between our protocol sampling polyA-enriched transcripts in the whole HCP versus their single-nucleic approach enriching for nucleic RNAs. In their study, increase in *Xist* did not reflect into measurable differences in the activity of the X chromosome, except for a few X-linked escape genes in a cell type-specific manner^36^. Taking advantage of our biological system, where we can discriminate the two X-chromosomes, we mined our RNAseq dataset resource to learn more about the endurance of XCI through aging. Our data revealed a skewing of XCI towards the BL6 chrX for 10 out of 11 mice (Fig. 6ii), consistent with previous studies^45^. This effect was not age-specific which contrast with the hematopoietic system for which aging-mediated enhanced skewing has been reported^77,33^. This may be explained by the differences in cell cycle turnover which is much higher during hematopoiesis than neurogenesis. Likewise, no signs of relaxation of XCI were observed in the aging HCP (Fig. 6D), which contrasts with the reported, albeit very subtle, alterations observed in the hematopoietic system^34,35^. While it is essential to acknowledge the potential for subtle effects on X-chromosome inactivation (XCI) stability with a larger sample size, our findings unequivocally indicate that XCI is remarkable resilience to physiological aging of the HCP.

In conclusion, our study represents a comprehensive investigation of the allele-specific DNA methylation and transcriptional landscape of the aging brain. While we focused on genomic imprinting and XCI, this dataset holds promise for exploring other monoallelic-specific phenomena that may be relevant to aging. We acknowledge certain limitations, including our focus on female and non-disease mice, the use of a reasonably small cohort of animals, and the lack of single-cell resolution. Nevertheless, our findings support the remarkable resilience of genomic imprinting and XCI in the face of epigenetic drift observed during healthy aging of the brain. The stability of these epigenetic processes with aging suggests their importance in preserving essential cellular functions throughout an entire individual’s lifespan.

## Acknowledgements

We would like to the members of the S.T.d.R.’s team for helpful discussions and the animal facility of the Instituto of Medicina Molecular for their help in maintaining the animal colonies. Work in S.T.d.R.’s team was supported by Fundação para a Ciência e Tecnologia (FCT) Ministério da Ciência, Tecnologia e Ensino Superior (MCTES), Portugal [IC&DT projects 2022.01532.PTDC and PTDC/BIA-MOL/29320/2017 as well as projects UIDB/04565/2020 and UIDP/04565/2020 of the Research Unit Institute from Bioengineering and Biosciences – iBB and LA/P/0140/2020 of the Associate Laboratory Institute for Health and Bioeconomy – i4HB]; S.T.d.R. and SM are supported by assistant research contracts from FCT/MCTES (2021.00660.CEECIND and CEECIND/02356.2021, respectively).

## Author contributions

S.T.d.R. conceived the study, supervised the project and together with M.E.-M. secured funding. S.M. maintained the BL6 and CAST mouse lines and sacrificed the mice, dissected brain areas, performed most of the molecular biology experiments, conducted IMPLICON experiments, analysed 5mC/5hmC measurements, IMPLICON and RNA-seq results. J.S. prepared the RNAseq alignment and analysed RNA-seq data under the supervision of M.E.-M.. D.O performed the 5mC/5hmC measurements. F.K. and M.E.-M. conducted bioinformatic analysis of IMPLICON. A.M. helped in RNAseq, IMPLICON and LC-MS/MS experiments. S.M. and S.T.d.R. wrote the manuscript with contributions of J.S. and M.E.-M.

## Competing interests

A.M. and F.K. are Altos Labs employees. The other authors declare no competing interests.

---

**Figure S1.**
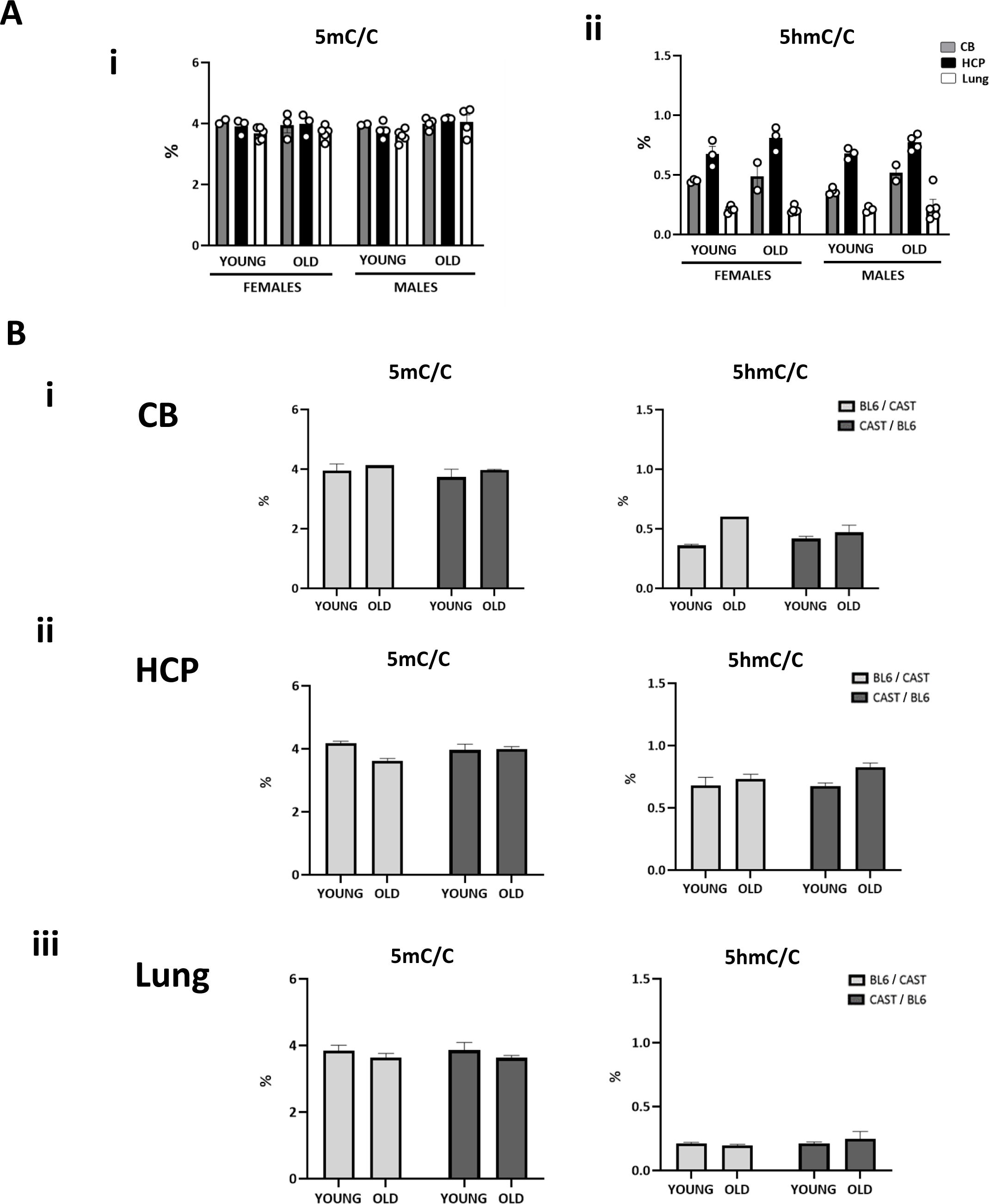
Evaluation of the influence of biological sex and the direction of cross on global DNA methylation level. **(A)** Influence of biological sex in 5mC/C **(i)** and 5hmC/C **(ii)** levels in cerebellum (CB), hippocampus (HCP) and lung of young and old females and males of F1 mouse hybrids of C57BL/6J (BL6) and Cast/EiJ (CAST) reciprocal crosses measured by Liquid Chromatography with tandem mass spectrometry (LC-MS/MS). Graphs represent the average percentage ± SEM of 5mC or 5hmC per total cytosines in young and old female (n=13) and male mice (n=9); (Female: CB - young n= 3, old n= 2; HCP - young n=3, old n=3; Lung - young = 5, old n= 5. Male: CB; young n= 4, old n= 2; HCP - young n=3, old n= 4; Lung - young n=4, old n= 5. Individual values are represented by dots. The data is the same as Fig. 2A, but separated by biological sex. **(B)** Influence of direction of the cross in 5mC/C and 5hmC/C levels in cerebellum (CB), **(i)**, hippocampus (HCP), **(ii)** and lung, **(iii)** of young and old of F1 mouse hybrids of BL6 and CAST reciprocal crosses measured by Liquid Chromatography with tandem mass spectrometry (LC-MS/MS). Graphs represent the average percentage ± SEM of 5mC or 5hmC per total cytosines in young and old BL6/CAST and CAST/BL6 F1 hybrid mice, (Female. CB young BL6/CAST n= 2, CAST/BL6 n=1, CB old BL6/CAST n=1, CAST/BL6 n=1; HCP young BL6/CAST n=2, CAST/BL6 n= 1; HCP old BL6/CAST n=1, CAST/BL6 n= 2; lung young BL6/CAST n=3, CAST/BL6 n= 2, lung old BL6/CAST n=3, CAST/BL6 n=2), (Male. CB young BL6/CAST n=2, CAST/BL6 n=2; CB old BL6/CAST n=, CAST/BL6 n= 2; HCP young BL6/CAST n= 1, CAST/BL6 n=2; HCP old BL6/CAST n= 2, CAST/BL6 n= 2; lung young BL6/CAST n=2, CAST/BL6 n= 2, lung old BL6/CAST n= 2, CAST/BL6 n=3). The data is the same as Fig. 2, but separated by direction of the cross.

**Figure S2.**
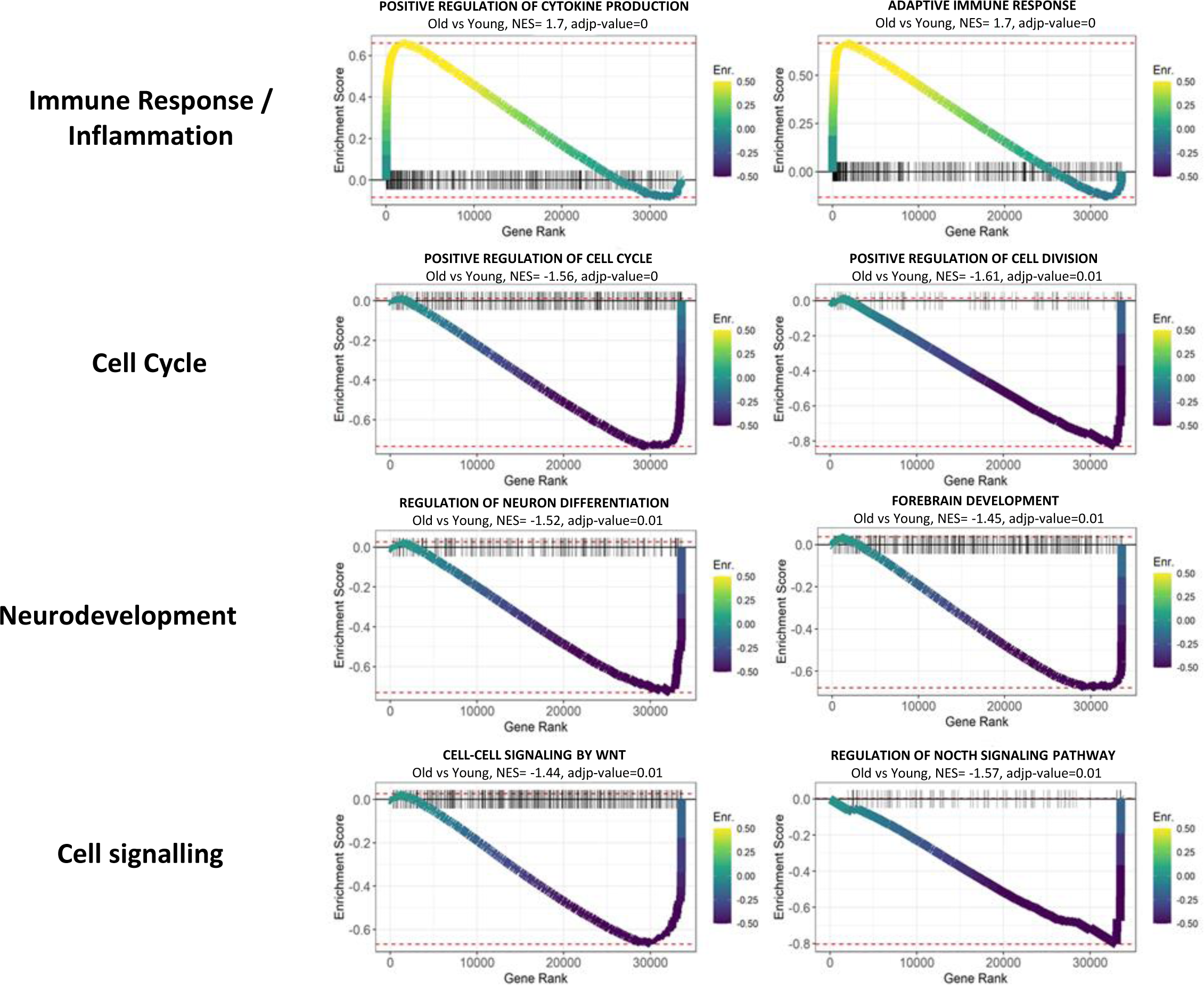
GSEA pathways differentially regulated in the HCP during aging. Running enrichment plots for selected top pathways enriched after GSEA analyses of differentially expressed genes in old vs young mice. For each plot the x-axis represents the ranking of differentially expressed genes by log2FC (old vs young mice) from upregulated (left) to downregulated (right) genes. Vertical lines indicate genes in the ranked list which were also represented in the pathway of interest. A running enrichment score is plotted on the y-axis which indicates the overall enrichment (NES > 0) or depletion (NES < 0) of the pathway in old vs young mice. The log adj p-value for each enriched gene set and normalized enrichment score (NES) are provided above each plot.

**Figure S3.**
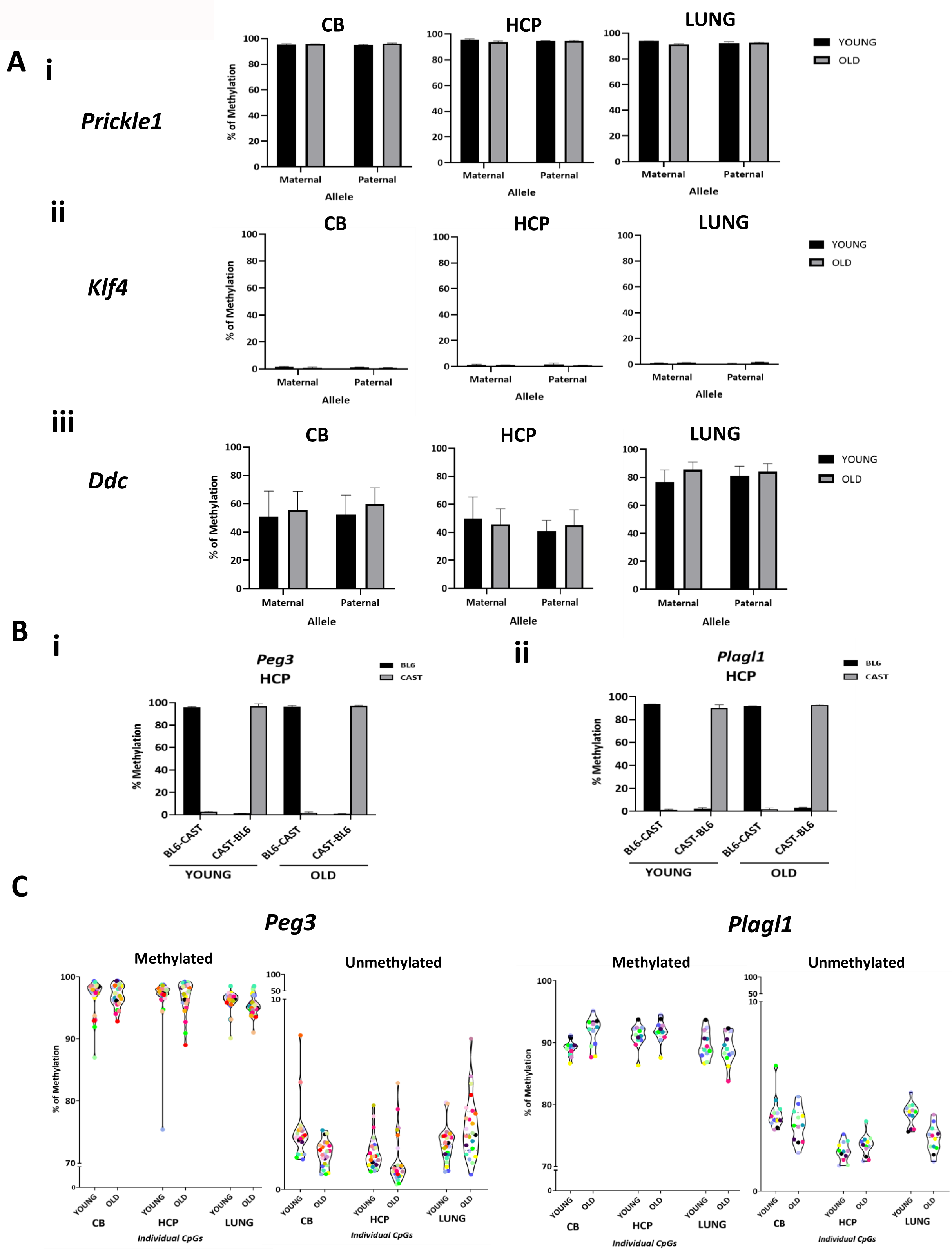
Methylation analysis of imprinted and control regions by IMPLICON. **(A)** Methylation analysis of *Prickle1* (methylated control) **(i)**, *Klf4* (unmethylated control) **(ii)** and *Dopa-decarboxylase* (*Ddc*) **(iii)** loci in cerebellum (CB), hippocampus (HCP) and lung of young and old females of F1 mouse hybrids of C57Bl/6J (BL6) and Cast/EiJ (CAST) reciprocal crosses. Barplots represents the mean ± SEM methylation levels measured at each CpG within each genomic region (n=4 for young and old female mice - (2 animals of each reciprocal cross per age group). **(B)** Influence of direction of cross in methylation levels for *Peg3,* **(i)**, and *Plagl1* ICRs, **(ii)**, in hippocampus (HCP) of young and old F1 mouse hybrids of BL6 and CAST reciprocal crosses measured by IMPLICON. Barplots represent the mean ± SEM methylation levels measured at each CpG within each genomic region (n=4 for young and old mice - 2 animals of each reciprocal cross per age group). **(C)** Violin plot displaying the mean of percentage of methylation of each individual CpGs for methylated and unmethylated alleles of the ICRs of the *Peg3* and *Plagl1* imprinted regions in cerebellum (CB), hippocampus (HCP) and lung of young and old female mice (n=4 for young and old mice - 2 animals of each reciprocal cross per age group).

**Figure S4.**
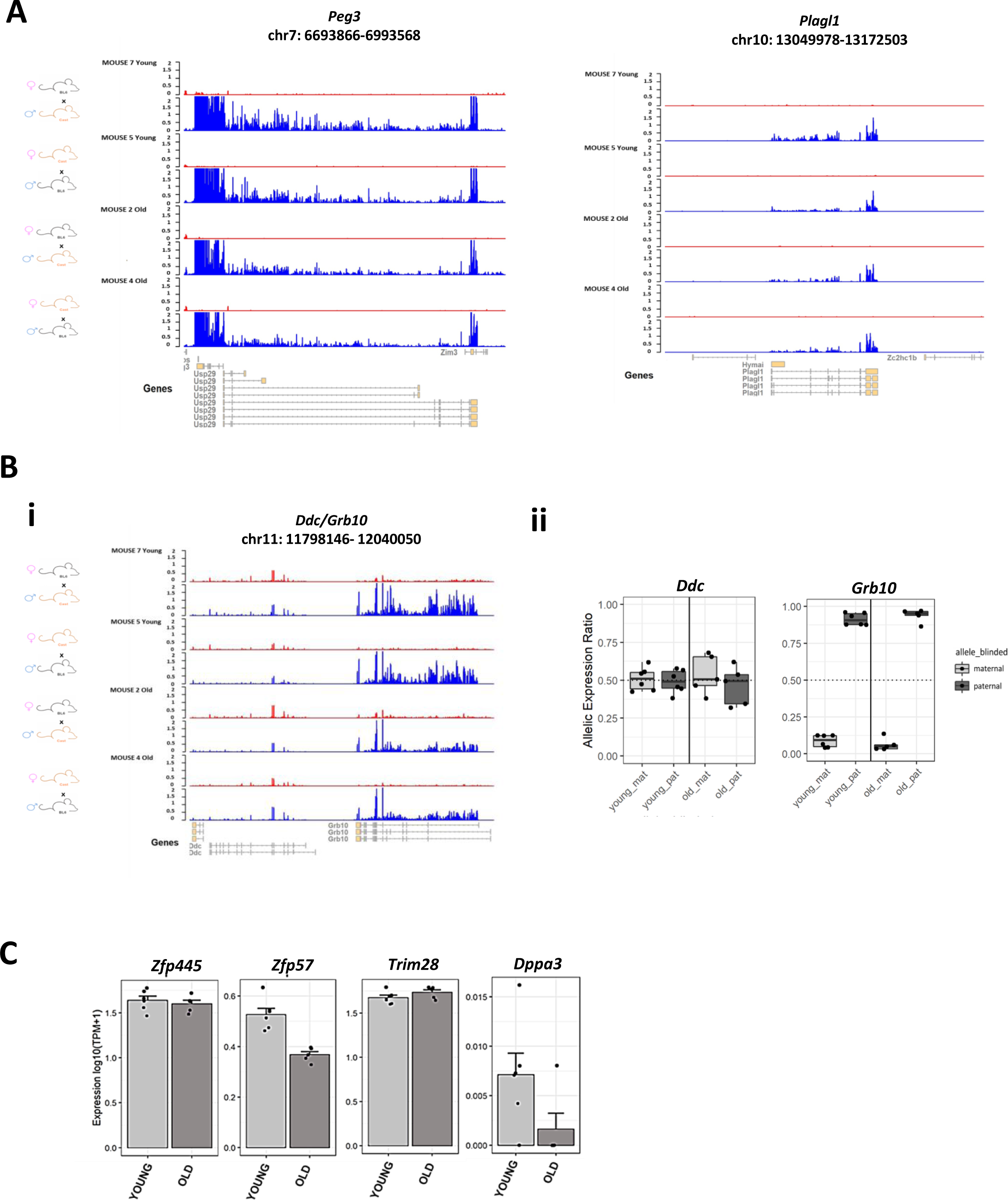
Stability of parental-of-origin allelic expression in HCP during aging. **(A)** RNA-seq allelic genome tracks showing the allelic expression of the imprinted genes *Peg3* and *Plagl1*. Two young and 2 old female mice from each reciprocal cross were chosen for display (red = maternal expression, blue = paternal expression). **(B)** RNA-seq allelic tracks of the *Ddc/Grb10* locus for the same mice **(i)**, and allele-specific expression of the *Ddc* and *Grb10* genes in the HCP of young (n=6) and old (n=5) mice measured by RNA-seq **(ii)**. **(C)** Bulk expression levels of genes involved in imprinting maintenance in young (n=6) and old (n=5) hippocampi in log10(TPM+1) (transcripts per million) measured by RNAseq.

**Figure S5.**
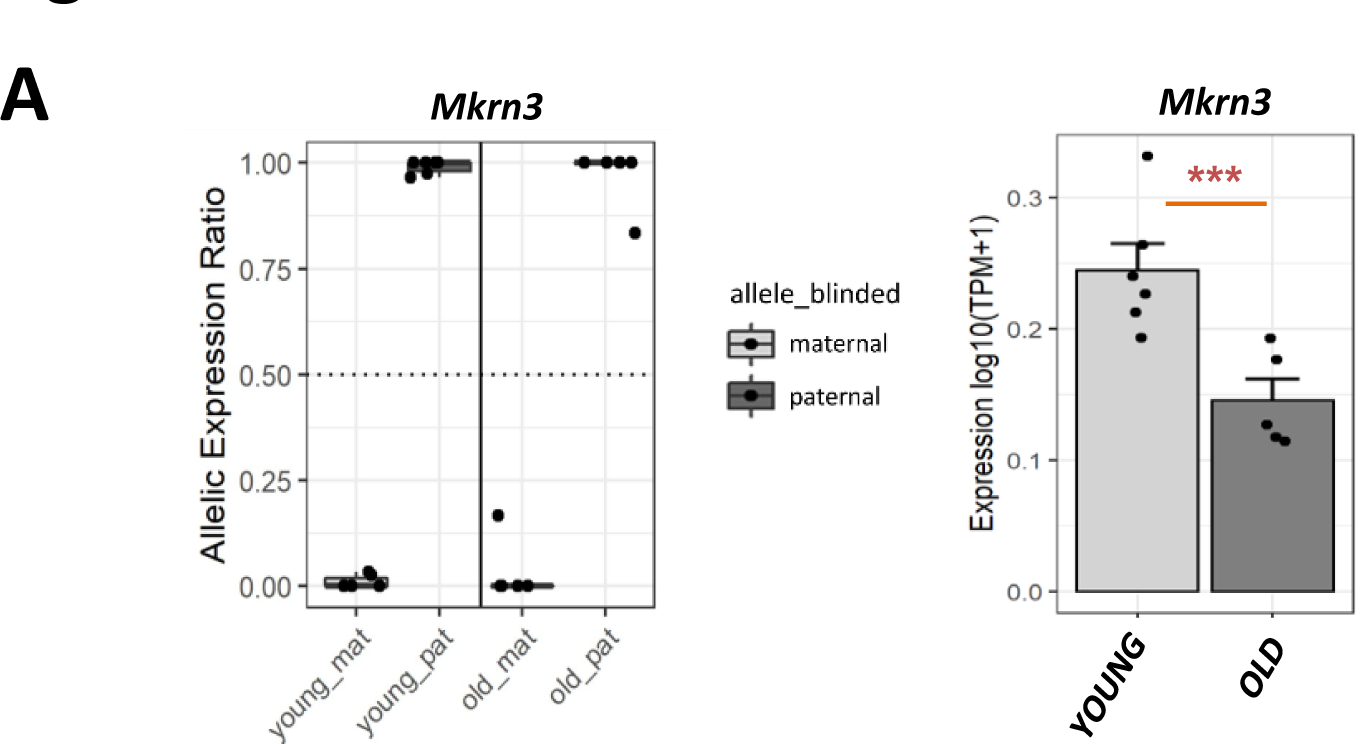
*Mkrn3* expression levels in young and old HCP. On the left, boxplot represents allelic expression ratio of maternal and paternal alleles of *Mkrn3* in young (n=5) and old HCP (n=6). On the right, barplot represents the average log10(TPM+1) expression of *Mkrn3* in young (n=6) and old HCP (n=5) (*P*adj < 0.05, log2FC < −1) statistically determined by DESeq2 (Wald test and BH-adjustment of p-values).

**Figure S6.**
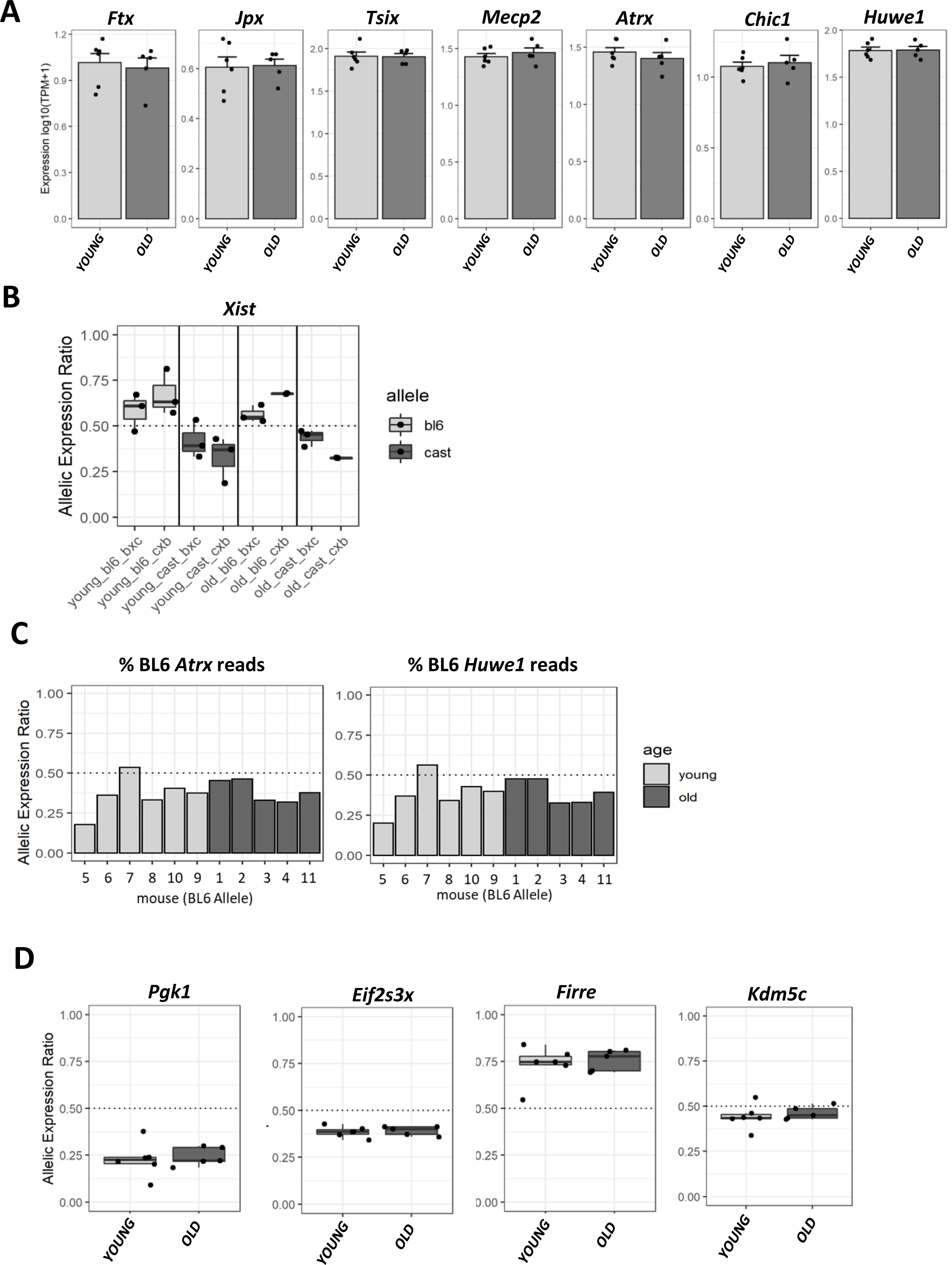
X-chromosome inactivation in young and old HCP. **(A)** Bar plots representing average bulk expression levels in log10(TPM+1) of noncoding genes on the X-inactivation center, *Ftx, Jpx* and *Tsix*, as well as *Mecp2*, *Chic1*, *Atrx* or *Huwe1* X-linked genes in young (n=6) and old HCP (n=5) measured by RNAseq. **(B)** Boxplots representing average allele-specific expression ratio of *Xist* reported as percentage of BL6 and CAST reads for young (n=6, of which 3 BL6/CAST and 3 CAST/BL6) and old mice (n=5, of which 3 BL6/CAST and 2 CAST/BL6) separated by the direction of the cross as measured by RNAseq. These data for percentage of BL6 reads is the same as shown in Fig. 6Bi but taking consideration the direction of the cross. Boxplot boundaries indicate the lower (1^st^) and upper (3^rd^) quartiles of allele expression ratios, with the middle line indicating the median value. **(C)** Barplots representing the percentage of allelic BL6 reads for the X-linked genes, *Atrx* and *Huwe1*, per each animal as measured by RNAseq. **(D)** Boxplots representing average allele-specific expression ratio reported as percentage of the BL6 reads for the escape genes *Eif2s3x* and *Kdm5c* in the HCP of young (n=6) and old (n=5) mice was measured by RNAseq. Boxplot boundaries indicate the lower (1^st^) and upper (3^rd^) quartiles of allele expression ratios, with the middle line indicating the median value.

